# Acquired resistance to PD-L1 inhibition is associated with an enhanced type I IFN-stimulated secretory program in tumor cells

**DOI:** 10.1101/2021.07.01.450417

**Authors:** Yuhao Shi, Melissa Dolan, Michalis Mastri, Amber Mckenery, James W. Hill, Adam Dommer, Sebastien Benzekry, Mark Long, Scott Abrams, Igor Puzanov, John M.L. Ebos

## Abstract

**Background:** Interferon (IFN) pathway activation in tumors can have dual, sometimes opposing, influences on immune responses. Therapeutic inhibition of programmed cell death ligand (PD-L1) – a treatment that reverses PD-1-mediated suppression of tumor-killing T-cells - is linked to alterations in IFN signaling; however, less is known about the role of IFNs after treatment resistance. Since IFN-regulated intracellular signaling can control extracellular secretory programs in tumors to modulate immunity, we examined the consequences of PD-L1 blockade on IFN-related secretory changes in preclinical models of acquired resistance.

**Methods:** Therapy-resistant cell variants were derived from orthotopically grown mouse tumors initially sensitive or insensitive to PD-L1 antibody treatment. Cells representing acquired resistance were analyzed for changes to IFN-regulated secretory machinery that could impact tumor progression.

**Results:** We identified a PD-L1 treatment-induced secretome (PTIS) that was enriched for several IFN-stimulated genes (ISGs) and significantly enhanced when stimulated by type I IFNs (IFNα or IFNβ). Secretory changes were specific to treatment-sensitive tumor models and found to suppress activation of T cells *ex vivo* while diminishing tumor cell cytotoxicity, revealing a tumor-intrinsic treatment adaptation with potentially broad tumor-extrinsic effects. When reimplanted *in vivo*, resistant tumor growth was slowed by the blockade of individual secreted PTIS components (such as IL6) and stopped altogether by a more generalized disruption of type I IFN signaling. *In vitro*, genetic or therapeutic methods to target PD-L1 could only partially recapitulate the IFN-enhanced PTIS phenotype, showing that *in vivo*-based systems with intact tumor:immune cell interactions are needed to faithfully mimic acquired resistance as it occurs in patients.

**Conclusions:** These results suggest that prolonged *in vivo* PD-L1 inhibition can ‘rewire’ type I IFN signaling to drive secretory programs that help protect tumors from immune cell attack and represent a targetable vulnerability to overcome acquired resistance in patients.

## Background

Cancer therapies can provoke unexpected (and often unwanted) cellular reactions that include the secretion of proteins such as growth factors and cytokines – many of which have been exploited as possible biomarkers of treatment effect or toxicity in patients^1–3^. Such therapy-induced secretomes (TIS) can also contribute to cancer progression, particularly in settings of acquired resistance where tumor cell populations adapt to treatments over prolonged periods^3^. For immune-checkpoint inhibitors (ICIs) that target the programmed cell death 1 (PD-1) pathway, early cytokine changes (e.g. IL6, IL8) in patients after treatment can correlate with initial responses^4,5^ or adverse events^6^, but less is known about whether tumor-specific secretory profile changes can be a cause or a consequence of acquired resistance^3,7^.

In this regard, tumoral control of secretory programs by interferons (IFNs) may be of interest in the setting of acquired resistance to PD-L1 inhibition for several reasons^3,8^. First, IFNs have been linked to multiple ICI treatment resistance mechanisms, mostly via the induction of IFN-stimulated genes (ISGs) activated by type I (α/β) and type II (γ) IFN subtypes. Currently the precise effect of IFNs on PD-L1 inhibitor efficacy remains enigmatic because they can, somewhat paradoxically, both protect and weaken immune defenses (often simultaneously)^9,10^. For instance, IFNs can boost antigen presentation (e.g. via beta-2-microglobulin and MHC-I expression) to improve PD-1 inhibitor responses ^11,12^, while also suppressing immune cell attack via the induction of T-cell inhibitory ligands^9,13^, NOS2^14^, and SerpinB9^15^, amongst many others^16,17^. Second, IFNs also can regulate a range of cellular processes that involve additional cytokine production^18,19^ that, in turn, can have positive and negative effects on tumor progression. Finally, several studies have now identified a crosstalk between tumor intrinsic PD-L1 functions and IFN signaling that control STAT3/Caspase 7 pathways^20^ and tumor cell DNA damage response^21^ – all of which can regulate protein production with extrinsic functions^22,23^. Currently it is unknown whether these IFN-controlled secretory programs are enhanced or inhibited in tumor cells in the context of acquired resistance where persistent PD-L1 blockade may impact immune-protective processes.

To examine this, we generated PD-L1 drug resistant (PDR) tumor cells to evaluate changes in secretory profiles regulated by IFN-signaling. PDR cell variants representing acquired resistance were generated using *in vivo* tumor models that were initially sensitive to anti-PD-L1 (αPD-L1) treatment and thus able to better mimic the complex tumor:immune cell interactions that occur during the progression to treatment failure. Using transcriptomic and proteomic analysis, we identified an αPD-L1 treatment-induced secretome (PTIS) signature in PDR cells that was enriched for ISGs and could be validated in multiple clinical and preclinical datasets involving αPD-L1 therapy. Type I IFNβ stimulation was found to potently enhance PTIS expression and IFN-controlled secretory products from PDR cells could shield tumors from CD8+ T cell cytotoxicity in immune cell co-culture studies. When reimplanted *in vivo*, PDR tumor growth slowed when IL6 (an ISG associated with PTIS) was inhibited and stopped altogether when type I IFN signaling was disrupted, suggesting IFN secretory changes represent a unique vulnerably in resistant cell populations. Interestingly, when therapeutic and genetic methods were used to mirror chronic PD-L1 inhibition *in vitro*, the IFN-enhanced PTIS effect was only partially recapitulated, indicating that tumor-intrinsic secretory changes depend, at least in part, on extrinsic host responses to treatment. Together, these results show that PD-L1 inhibition disrupts immune protective secretory programs in treatment-sensitive tumors and identify a unique ‘rewiring’ of type I IFN signaling that could improve outcomes if therapeutically targeted in patients with acquired resistance.

## Results

### Acquired resistance to PD-L1 inhibition increases secretory profiles enriched for type I IFN regulated genes

To examine acquired resistance to PD-L1 inhibition, the PD-1 pathway *inhibitor-sensitive* murine breast tumor EMT6 cell line ^24–26^ was implanted orthotopically in BALB/c mice and treated with αPD-L1 (clone 80) or IgG control antibody **(Fig 1a; schematic shown).** Following continuous treatment, a PD-L1 drug-resistant (PDR) cell variant (EMT6-PDR) was selected from mice with tumors that resumed growth after an initial significant delay (**Fig 1b; circles shown**). Transcriptome RNA-sequencing of EMT6-PDR and EMT-P (parental) tumor tissues revealed multiple genes to be up- or down-regulated **(Fig 1c)**. Gene-set enrichment analysis (GSEA) showed EMT6-PDR tumors to be significantly enriched for genes associated with extracellular matrix, growth factor, and cytokine signaling pathways, several of which were secreted and IFN-regulated (**Fig 1d**). Using the Gene Ontology (GO) database term GO:0005576 consisting of products outside or unattached to the cell ^8^, secretory genes were found to increase in EMT6-PDR tumor transcripts and associate with inflammatory signaling, wound healing, and immune cell function/migration **(Fig 1e).** Since many of these processes also associate with IFN signaling ^19,27^, we examined IFN-regulated genes using the Interferome database - a compilation of published *in vitro* and *in vivo* experimental datasets identifying transcriptomic and proteomic changes after IFN treatment ^28^. Compared to P controls, EMT6-PDR tumors had several IFN-related genes up- and down-regulated (54% and 63%, respectively), with type I IFN gene upregulation the most common (22% of total) (**Fig 1f)**. To confirm association of IFN regulated genes in EMT6-PDR tumors, we assessed for relative enrichment of IFN signaling gene sets found in several publications ^9,10,29–31^ and in the Hallmark Molecular Signatures Database (MSigDB) ^32^ where positive enrichment was observed for all datasets examined (**Table S1**; **Fig 1g).** Similar positive enrichment was observed when GSEA was performed on an IFNγ-associated gene-set identified in durvalumab-treated nonsmall cell lung carcinoma (NSCLC) patient tumor biopsies ^33^ **(Fig 1h)**. At the individual gene level, qRT-PCR analysis confirmed upregulation of several ISGs and type I IFNs in EMT6-PDR cell variants **(Fig 1i)**. To test whether this was specific to acquired resistance, we examined IFN and ISG expression in PDR tumors derived from an innately resistant model. To do this, we first implanted the PD-1 pathway inhibitor-*insensitive* murine kidney tumor RENCA cell line orthotopically into BALB/c mice^34^ and, following treatment with αPD-L1 or IgG antibody (**Fig 1j; schematic shown**), we then selected a RENCA-PDR tumor cell variant that did not respond to treatment (**Fig 1k; circles shown**). GSEA of IFN-associated gene sets showed a negative enrichment in RENCA-PDR cells (**Fig 1l)**. Taken together, these results demonstrate that αPD-L1 treatment can induce ISG-related secretory gene changes in tumors that are enhanced in acquired resistance settings.

**Figure 1.**
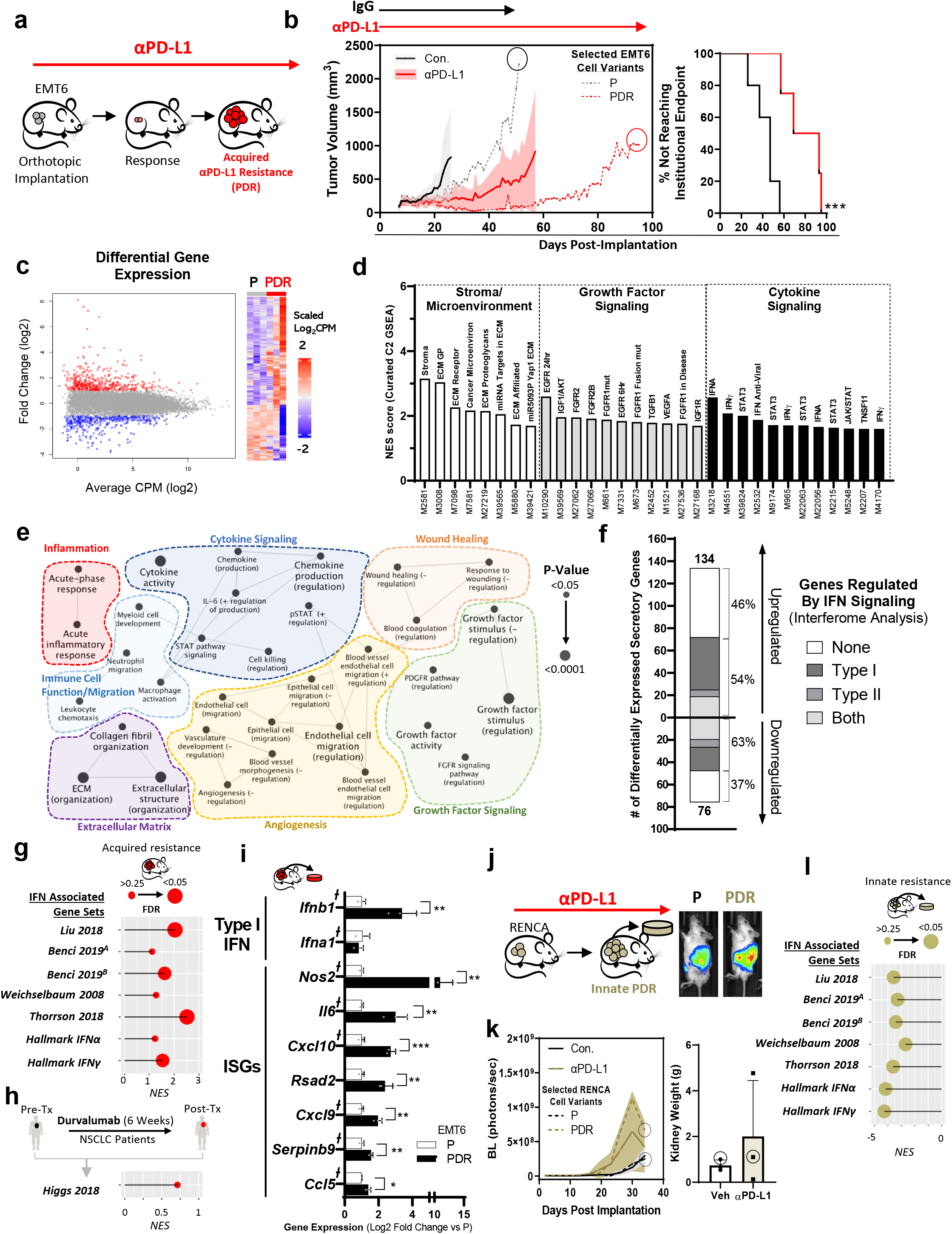
Acquired anti-PD-Ll drug resistance (PDR) increases secretory profiles enriched for type I IFN regulated genes. **(a)** Schematic showing orthotopic breast EMT6 model of acquired resistance to αPD-L1 inhibition **(b)** Continuous αPD-L1 treatment in BALB/c mice (n=3-4) bearing orthotopically-implanted mouse mammary EMT6 cells (left) and the time taken to reach endpoint defined by primary tumor size (2000m^3^) and animal morbidity (right). **(c-f)** RNA sequencing analysis of EMT6-P and EMT6-PDR tumor tissues. **c)** Differentially expressed genes (Log_2_ [Fold Change] ≤ −2 or ≥ 2) in EMT6-PDR as summarized by dot plot (left) and heatmap (right). Red = upregulated; blue = downregulated. **d)** Summary of stroma/tumor microenvironment, growth factor signaling, and cytokine signaling gene sets with significant positive enrichment found in PDR tumors via GSEA of all canonical pathways (C2, Molecular Signatures Database Collection). **(e)** Cytoscape GO analysis of significantly enriched biological processes in upregulated secretory genes, grouped by signaling categories. Size of circles correspond to FDR significance of each process and lines represent term-term interactions defined by Kappa score. **(f)** Bar graph representing Interferome Database secretory genes up- and down-regulated in EMT6-PDR (compared to EMT6-P). **(g-h)** GSEA of EMT6-PDR tumors (compared to P controls) showing lollipop plot representing **(g)** NES of published/Hallmark IFN gene sets, and **(h),** NES of an IFN-specific gene-set identified in αPD-L1-treated (durvalumab) NSCLC patients (described in Ref 33). Size of circles correspond to FDR significance. See Table S1 for details. **(i)** Type I IFNs and additional ISGs in EMT6-PDR selected cell variants (qRTPCR). *┼* represent genes associated with secretory proteins. **(j)** Schematic showing orthotopic kidney RENCA tumor model of innate resistance to αPD-L1 inhibition (left) and BLI of selected tumor variants on day of endpoint (right). **(k)** BLI quantification of murine RENCA orthotopic tumor growth (left) and kidney weight at endpoint (right) with continuous αPD-L1 treatment (Balb/c mice; n =3). **(l)** GSEA of RENCA-PDR tumor cells (compared to P controls) showing lollipop plot representing NES of published/Hallmark IFN gene sets. Size of circles correspond to FDR significance. See Table S1 for details. Parental (P); αPD-L1 Drug Resistant (PDR); Control (Con); Gene Set Enrichment Analysis (GSEA); Gene Ontology (GO); interferon stimulated genes (ISGs); Counts per million (CPM); Bioluminescence (BLI); normalized enrichment scores (NES); Non-small cell lung carcinoma (NSCLC). αPD-L1 (clone 80) and IgG/PBS were administered at 250μg/mouse every 3 days until endpoint. Primary tumor burden was assessed by caliper measurement. Time to institutional endpoint was assessed by Kaplan-Meier. PDR cell variant tumor growth shown as dotted line and time of tumor selection shown with circle for 1B and 1I. Selected EMT6-PDR and EMT6-P variants were maintained in vitro with respective αPD-L1 or IgG antibody (see Methods for details). Quantitative data shown as mean ± SD. * p<0.05, ** p<0.01, *** p<0.001, ****p<0.0001

### A PD-L1 treatment-induced secretome (PTIS) is enriched in PD-L1 treatment-sensitive tumors

We next tested whether IFN-enriched secretory effects could be confirmed in other tumor models reported to be initially PD-L1 treatment-sensitive. To do this, we developed a composite of secretory genes and proteins found to be increased in EMT6-PDR tumors based on RNAseq, qRT-PCR, cytokine array, and ELISA analysis (**See Table S2; Fig S1; Methods**). From this, we identified a αPD-L1 treatment-induced secretome (PTIS) signature consisting of 12 up-regulated molecules, all representing IFN-regulated genes identified via the Interferome Database (**Fig 2a**). Enrichment for the PTIS signature was tested using publicly available NCBI GEO and dbGAP whole transcriptome datasets from published preclinical and clinical studies involving αPD-L1 treated tumors. In 5 preclinical studies examined, 3 were reported as αPD-L1 *treatment-sensitive* (Lan et al ^24^; Sceneay et al ^35^; Efremova et al ^36^) and 2 were αPD-L1 treatment-*insensitive* (Sceneay et al ^35^ and RENCA-PDR tumor cells from current study) (**See Methods**). PTIS signature expression was increased in all αPD-L1 treatment-*sensitive* models as defined by average counts per million (CPM) levels, with 2 of 3 models demonstrating significant positive GSEA enrichment and 1 of 3 models showing significance by both CPM expression and GSEA enrichment (**Fig 2b**). Conversely, PTIS signature expression was decreased in αPD-L1 treatment-*insensitive* models (**Fig 2c**). In 2 clinical studies examined, tumor biopsies were taken from non-small cell lung carcinoma (NSCLC) (Gettinger et al. ^37^) and Merkel cell carcinoma (MCC) (Paulson et al. ^38^) patients reported to be initially sensitive to αPD-L1 treatment (**See Methods**). In the NSCLC samples, the PTIS signature expression increased with significant positive GSEA enrichment in bulk RNAseq data from patients who developed acquired resistance **(Fig 2d)**. In MCC samples, single-cell RNAseq datasets also showed increased PTIS signature expression in tumor, macrophage, and T-cell enriched cell compartments after clustered analysis **(Fig 2e; See Methods)**. Notably, we generated a separate PTIS using only genes *downregulated* in EMT6-PDR cells (termed ‘PTIS^DOWN^’) and found dataset validations not consistent, suggesting *upregulated* PTIS genes are more representative of acquired resistance **(Fig S2; See Supplemental Results)**. Together, these findings demonstrate that the IFN-enriched PTIS is enriched in multiple preclinical/clinical tumors initially *sensitive* to αPD-L1 treatment and can occur independent of cancer type.

**Figure 2:**
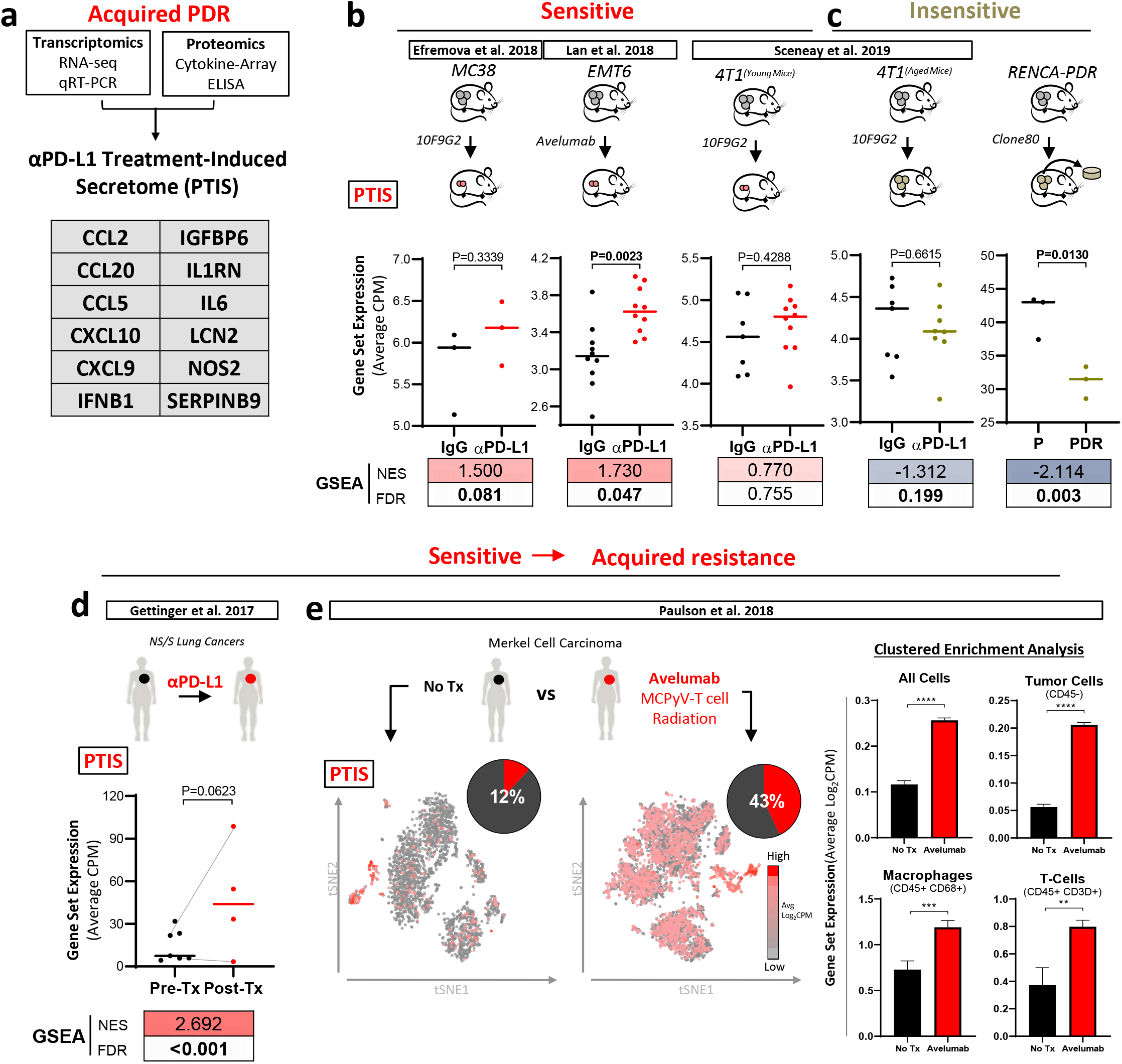
An upregulated PTIS signature is enriched in clinical and preclinical models sensitive to αPD-L1 treatment. **(a)** Generation of an αPD-L1 treatment-induced secretome (PTIS) comprised 12 upregulated genes identified from transcriptome and proteomic analysis using EMT6-PDR cells. **(b-e)** PTIS expression in published bulk and single cell RNAseq datasets involving αPD-L1 treatment in preclinical and clinical studies. RENCA-PDR model from this study included. **(b-c)** Preclinical studies: PTIS expression using average CPM expression and GSEA in datasets taken from tumor models involving αPD-L1 treatment and found to be **b)***treatment-sensitive* (GEO: GSE130472, GSE93017, GSE107801) or **c)** treatment-*insensitive* (GEO: GSE130472; RENCA-PDR). Data is compared to vehicle/IgG-treated controls. **(d-e)** Clinical studies: PTIS expression using average CPM expression and GSEA in datasets taken from tumor biopsies of αPD-L1 treatment-sensitive patients. **d)** NSCLC patients (dbGAP # phs001464.v1.p1): bulk RNAseq from Pre-Tx and Post-Tx tumor sample comparisons (Gray lines indicate matched Pre- and Post-tx samples). **e)** MCC patients (GEO: GSE118056): single-cell RNAseq from untreated (No-Tx) or treated (avelumab, MCPyV-T cell, radiation) tumor samples with Tsne plots (left) representing average log_2_ CPM expression of PTIS in whole dataset, and bar graphs (right) representing clustered enrichment analysis populations identified by markers for tumors (CD45-), macrophages (CD68+), and T cells (CD3D+). Tumor sample that received No-Tx was compared to treated. αPD-L1 Treatment-Induced Secretome (PTIS); αPD-L1 Drug Resistant (PDR);Counts per million (CPM); Gene set enrichment analysis (GSEA); False Discovery Rate (FDR); Gene Expression Omnibus (GEO); GEO Series records (GSE); database of Genotypes and Phenotypes (dbGaP); t-distributed stochastic neighbor embedding (tsne); Treatment (Tx); non-small cell lung carcinoma (NSCLC); Merkel cell carcinoma (MCC). #Indicates genes not part of the Interferome database. Significance represented as * p<0.05, ** p<0.01, *** p<0.001, **** p<0.0001. Bolded numbers for GSEA represent FDR<0.25 (see Methods).

### Type I IFN stimulation enhances PTIS expression in PDR cells

ISGs found in the PTIS represent secreted factors that have diverse and at times opposing effects on tumor development and growth^18,39^. For instance, cytokines such as IL6, CXCL9, CXCL10 can promote and inhibit anti-tumor immunity^40–42^, regulate tumor angiogenesis^43^, and have autocrine signaling in tumor cells^44–46^. Additionally, extracellular (but non-secreted) ISGs such as antigen presentation molecules MHC-I^47^ and PD-L1^48^ can also affect how tumor cells respond to ICI treatment, often with opposing effects on immune activation. Since secreted and non-secreted ISGs can be regulated by type I IFNs (α, β) binding to IFN alpha receptor (IFNAR) ^49^, we tested PDR cell variants after stimulation with recombinant type I IFNs (α or β) or type II IFN (γ). While multiple PTIS genes *(IL6, Nos2, Serpinb9)* were upregulated by type I IFN stimulation, this effect was uniquely enhanced in EMT6-PDR cells (**Fig 3a)**. *IL6* gene expression was the most robustly increased by type I IFN stimulation in EMT6-PDR cells (**Fig 3b; heatmap summarizing relative expression shown**), which we confirmed at the protein level with significant enhancement of secreted IL6 found in EMT6-PDR cell conditioned media (CM) (**Fig 3c**). Control experiments to test whether known anti-proliferative effects of IFN stimulation ^50^ could account for PTIS expression were ruled out as EMT6-P and PDR cells did not consistently respond differently to IFN exposure (**Fig S3**). Next, to test whether the IFN-enhanced PTIS was primarily the result of type I IFN signaling regulation, IFNAR1 was knocked down in EMT6-P and PDR cells (designated ‘IFNAR1^KD^’) via short hairpin RNA **(Fig 3d**). Our results showed that IFNβ-stimulated increases in IL6 expression could be reversed in EMT6-PDR-IFNAR1^KD^ cells **(Fig 3e)**. To further connect the PTIS and IFN-regulated programs, our next studies examined intracellular STAT proteins known to be activated by IFN-signaling **(Fig 3f; blotting replicates in Appendix 1)**. In EMT6-PDR cells, total STAT1 expression was decreased compared to P controls while total STAT3 levels remained unchanged **(Fig S4a-b; densitometry analysis)**. Yet, when treated with IFNβ, both pSTAT1 and pSTAT3 had significant increases in EMT6-PDR cells compared to P controls, an effect that was partially reversed in PDR-IFNAR1^KD^ cells **(Fig 3g-h; densitometry analysis)**. Confirmatory studies were performed for pSTAT3 levels using ELISA assays **(Fig S4c)**. Taken together, these results suggest intracellular pSTAT3 signaling is uniquely enhanced in EMT6-PDR cells following IFNβ stimulation. Next, we assessed immune-regulating non-secreted cell surface ISGs PD-L1 and MHC-I. Interestingly, and in contrast to the enhanced levels of secretory ISGs observed, PD-L1 **(Fig 3i)** and MHC-I **(Fig 3j)** were found to be decreased in PDR cells at baseline and after IFNβ stimulation compared to P controls. Overall, these results suggest that type I IFN stimulation uniquely enhances secretory programs after acquired resistance, including robust increases in IL6 expression and STAT signaling; but paradoxically, decreases immune-modulating ISGs expressed on the cell surface in the same setting (**Fig 3k summary and Fig S4d statistics summary)**. This raises the question of whether these secretory and cellular (non-secreted) proteins, known to have opposing roles in immunity, might influence (or be influenced by) immune cell populations that are part of the anti-tumor response.

**Figure 3:**
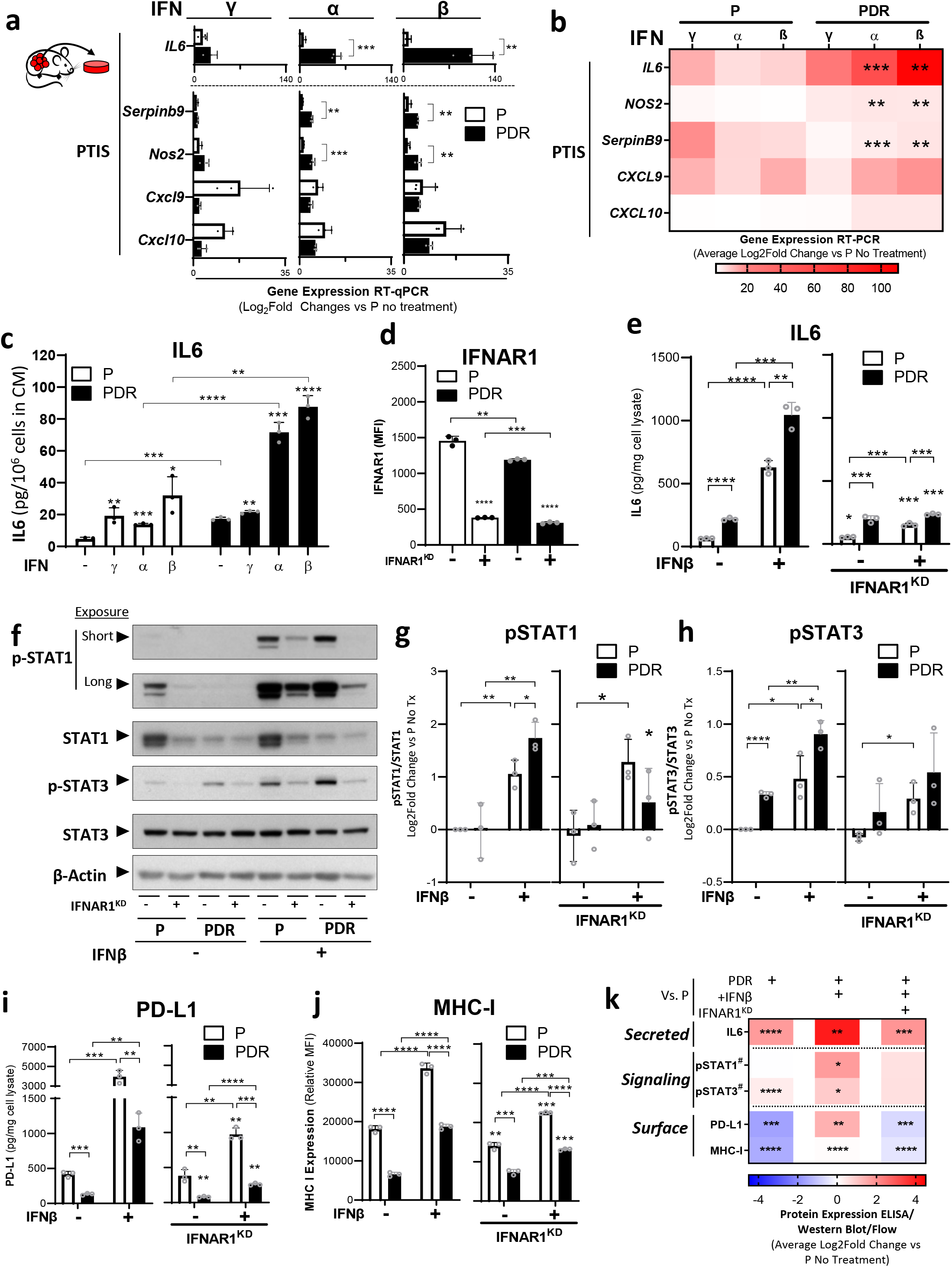
Type I IFN stimulation enhances PTIS after acquired PD-L1 resistance. **(a)** ISGs after stimulation with type I/II IFNs in EMT6-PDR cells shown as relative to untreated P controls and represented as bar graphs (qRT-PCR). **(b)** Heatmap summary of results in (a). Statistics comparing PDR versus P for each condition. **(c)** Secreted IL6 levels in EMT6-P and -PDR tumor cell CM after IFNγ, α, and β stimulation (ELISA). **(d)** IFNAR1 expression in EMT6-P and -PDR before and after knockdown of IFNAR1 and respective vector controls (flow cytometry). **(e)** IL6 expression in lysates of EMT6-P and -PDR before and after knockdown of IFNAR1^KD^, and after IFNβ stimulation (ELISA). **(f)** Phosphorylated and total levels of STAT1/3 in lysates of EMT6-P and -PDR before and after knockdown of IFNAR1^KD^ following IFNβ stimulation (Western Blot). **(g-h)** Densitometry quantification of western blots shown in (f) representing **(g)** relative phosphorylated STAT1 compared to total STAT1 and **(h)** phosphorylated STAT3 levels compared to total STAT3 levels. **(i)** PD-L1 expression in lysates of EMT6-P and -PDR before and after knockdown of IFNAR1^KD^, and after IFNβ stimulation (ELISA). **(j)** MHC-I expression in EMT6-P and -PDR before and after knockdown of IFNAR1^KD^, and after IFNβ stimulation (flow cytometry). **(k)** Heatmap summary of protein expression analysis (e-j). Statistics comparing PDR versus P for each condition. Parental (P); PD-L1 Drug Resistant (PDR); Conditioned Media (CM); Mean Fluorescent Intensity (MFI); IFN stimulated genes (ISGs); IFNAR1 knockdown (IFNAR1^KD^); shRNA vector control (shCon; shown here as a ‘-’); Cells were treated with 10ng/ml of IFNs and collected after 15mins (western blot shown in f), and 5 days (IL6, PD-L1, MHC-I). * p<0.05, ** p<0.01, *** p<0.001, **** p<0.0001 indicate significance compared untreated controls unless otherwise shown (lines). #STAT activation measured by comparing phosphorylated STAT protein compared to total STAT protein levels.

### Immune-suppression and -stimulation by PDR cells is IFN-regulated

To test this, we first performed CIBERSORT/ImmuCC tissue deconvolution analysis to identify immune cell populations in PDR tumors using mouse-specific gene scores in RNAseq data (described in ^51,52^). EMT6-PDR tumors had higher total immune scores **(Fig 4a)**, with activated cytotoxic CD8+ T lymphocyte (CTL) and M2 macrophage scores significantly decreased **(Fig 4b)**, indicating that anti-tumor immune responses may be suppressed after acquired PD-L1 resistance. To examine this directly, we next tested EMT6-P/PDR cells for cell cytotoxicity in co-culture studies with dissociated mouse splenocytes containing αCD3/αCD28 activated CD8+ T cells measured by flow cytometry (**Fig 4c; representative images shown**). We found tumor markers for apoptosis (annexin V) and cell death (7-AAD) to be significantly decreased in EMT6-PDR cells compared to P controls (**Fig 4d)**, suggesting PDR cells have an underlying immune-protective effect. Notably, this protection significantly decreased in PDR cells when IFNAR1 expression was knocked down (**Fig 4e; relative comparisons shown).** To test whether secretory factors released from PDR cells could influence CTL functions, we next measured CD8+ T-cell proliferation^53^, CD69 (an early T-cell activation marker^54^), and the T-cell effector proteins Granzyme B (GRMB) and IFNγ because they are capable of cell-killing^55,56^ (**Fig 4f and Fig S5a; schematics shown**). Conditioned media (CM) from EMT6-PDR cells significantly decreased T-cell proliferation (**Fig 4g, shown as % divided)**, CD69 **(Fig 4h)**, IFNγ **(Fig S5b)**, and GRMB **(Fig S5c).** Interestingly, these decreases could be significantly weakened (reversed) when CM from EMT6-PDR-IFNAR1^KD^ cells were used in the same experimental setup, suggesting the secretory products in PDR cells have an overall immune-suppressive effect that is IFN-regulated (**Figs 4g-h; Figs S5b-d**). Next, we conducted identical experiments but, rather than using CM, we combined tumor and splenocyte cells together in co-culture, with the goal to simultaneously examine both contact*independent* (secretory) and *contact-dependent* cellular interactions **(Fig 4i and Fig S5e; schematics shown)**. Our results show that, instead of the decreased immune cell activity seen with CM, co-culture produced significant increases in T-cell proliferation **(Fig 4j)** and CD69 (**Fig 4k),** or no changes in IFNγ **(Fig S5f)** and GRMB (**Fig S5g**). This indicates that contact-dependent tumor:immune cell effects can offset (and even reverse) contact-independent immune-suppression. Identical co-culture studies performed with EMT6-PDR-IFNAR1^KD^ cells significantly boosted some (but not all) immune-cell activation markers even further with T-cell proliferation and CD69 expression increasing (**Figs 4j-h)**, while GRMB or IFNγ remained unchanged **(Figs S5f-h**). A heatmap summary of contact-independent (secretory) and contact-dependent co-culture studies are detailed in **Fig 4l**. Together, these results show acquired resistance to PD-L1 inhibition ‘rewires’ tumoral IFN-signaling to produce *immune-suppressive* secretory changes and anti-apoptotic limiting stimuli; while simultaneously producing an opposing *immune-stimulatory* effect via contact dependent processes (**Fig 4m**).

**Figure 4:**
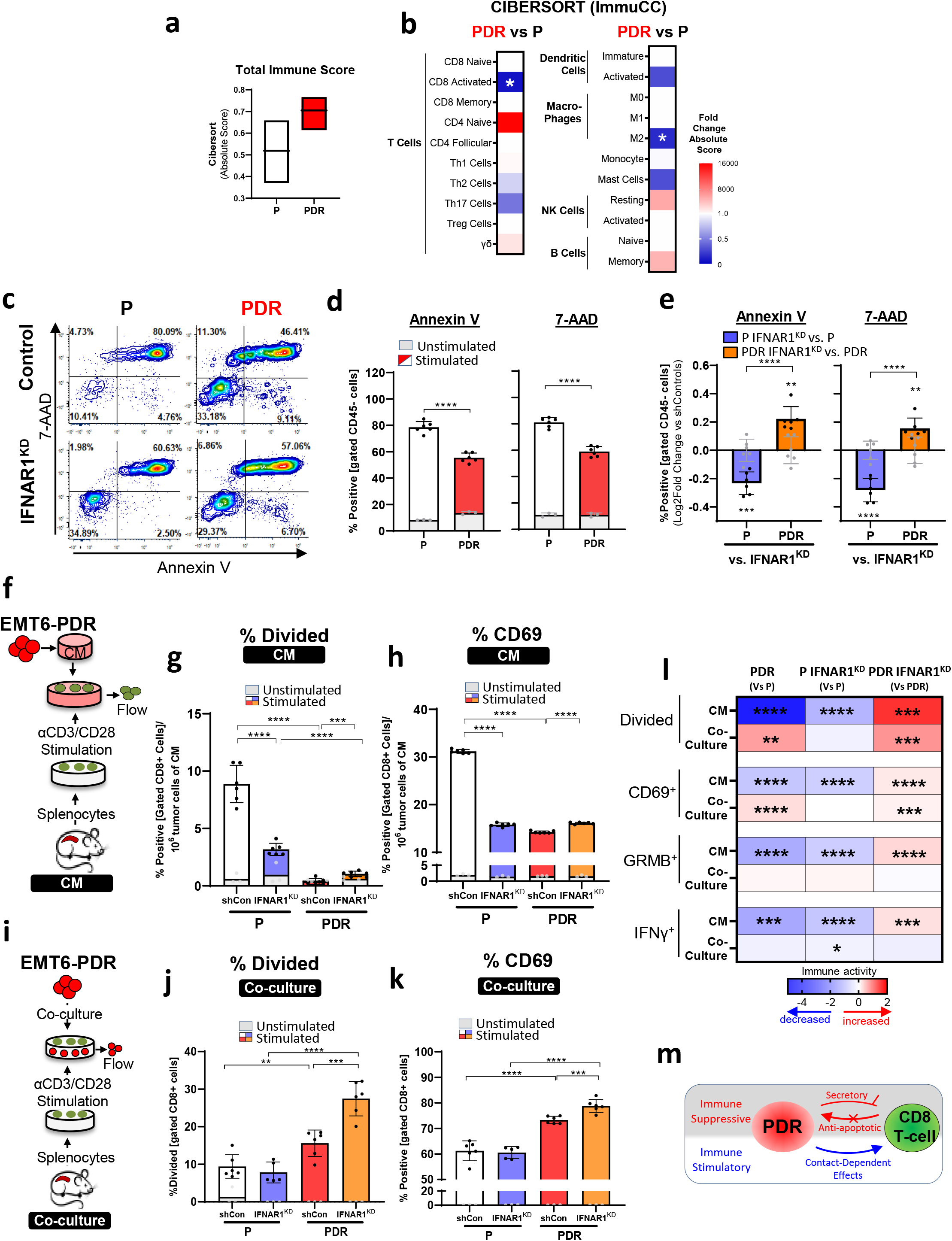
PDR immune-protective secretory changes are IFN signaling-dependent. **(a)** Cibersort tissue deconvolution analysis of EMT6-P and -PDR RNAseq data using ImmuCC mouse signature with box-plot representing absolute total immune score. **(b)** Heatmap representing log2 fold change of absolute scores of various immune signatures of results from (a). **(c-e)** Apoptosis (Annexin V) and cell death (7-AAD) staining of CD45-gated tumor cells after co-incubation with splenocytes showing **(c)** representative contour plots of stimulated splenocyte groups, **(d)** EMT6-P and –PDR variants, and **(e)** EMT6-P and –PDR-IFNAR1^KD^ variants compared to controls (flow cytometry). **(f)** Schematic of BALB/c–derived splenocyte proliferation and activation following incubation EMT6-P and –PDR CM for experiments in g-h. **(g)** CD8+ splenocyte division (CSFE dilution) after co-incubation with CM derived from EMT6-P and – PDR control and respective IFNAR1^KD^ variants **(h)** CD8+ splenocyte activation marker expression (CD69) after co-incubation with CM derived from EMT6-P and –PDR control and respective IFNAR1^KD^ variants. **(i)** Schematic of Balb/c–derived splenocyte proliferation and activation following co-culture with EMT6-P and –PDR cells for experiments in j-k. **(j)** CD8+ splenocyte division (CSFE dilution) after co-culture with EMT6-P and –PDR control and respective IFNAR1^KD^ variants. **(k)** CD8+ splenocyte activation marker expression (CD69) after co-culture with EMT6-P and –PDR control and respective IFNAR1^KD^ variants. **(l)** Heatmap summary of results from CM and co-culture experiments. **(m)** Schematic summary of immune suppressive and stimulatory effects of PDR cells. Parental (P); PD-L1 Drug Resistant (PDR); IFNAR1 knockdown (IFNAR1^KD^); shRNA vector control (shCon); Conditioned Media (CM) * p<0.05, ** p<0.01, *** p<0.001, ****p<0.0001 compared to vector controls unless noted otherwise. For C, F, H, and L white bars represent vector controls

### Inhibition of PTIS regulators selectively inhibits PDR tumor growth

To examine whether tumor growth may be altered by these opposing immune-modulating effects *in vivo*, we evaluated the impact of PTIS blockade *indirectly* by disrupting IFN-regulation or *directly* by blocking one secretory factor, such as IL6. To do this, we first implanted EMT6-P and EMT6-PDR cells orthotopically in BALB/c mice and found tumors grew at similar rates, suggesting immune-suppressive/promoting functions in PDR cells may offset to yield minimal changes in tumor growth kinetics. However, when type I IFN signaling was disrupted in the same PDR cells, we observed markedly opposing effects on tumor growth (**Fig 5a, AUC analysis shown in inset)**. IFNAR1^KD^ in EMT6-P tumors significantly *increased* tumor growth, whereas IFNAR1^KD^ in EMT6-PDR tumors significantly *decreased* tumor growth, compared to respective controls (**Fig 5b, AUC analysis shown in inset)**. While growth promotion by type I IFN signaling blockade in treatment-naïve tumors has been reported previously^57,58^, our results suggest PDR tumors have a unique vulnerability to IFN-signaling, perhaps the result of reducing the immune protective effect of the PTIS. To examine this further, anti-mouse IL6 (α-mIL6) antibody was administered to BALB/c mice bearing orthotopically-grown EMT6-P and EMT6-PDR tumors. PDR tumor variants treated with α-mIL6 showed significant tumor inhibition compared to P controls, but this inhibition was more muted than the more generalized PTIS blockade observed when IFN-signaling was disrupted (**Fig 5c, AUC analysis shown in inset; Fig 5d, comparisons shown to respective controls)**. Interestingly, PTIS-related vulnerabilities in PDR tumor growth were found to extend to metastasis with post-mortem analysis at experiment endpoint showing enhanced inhibition of tumor skin/abdominal wall invasion in PDR tumor variants after IFNAR1^KD^ and IL6 inhibition compared to P controls **(Fig 5e)**. Taken together, these results show that targeting PTIS has minimal or even growth promoting effects in untreated tumors yet, after acquired resistance to PD-L1 inhibition, tumors have a unique IFN-regulated vulnerability linked to PTIS expression that, when targeted directly or indirectly, has enhanced anti-tumor potency (**Fig 5f**).

**Figure 5.**
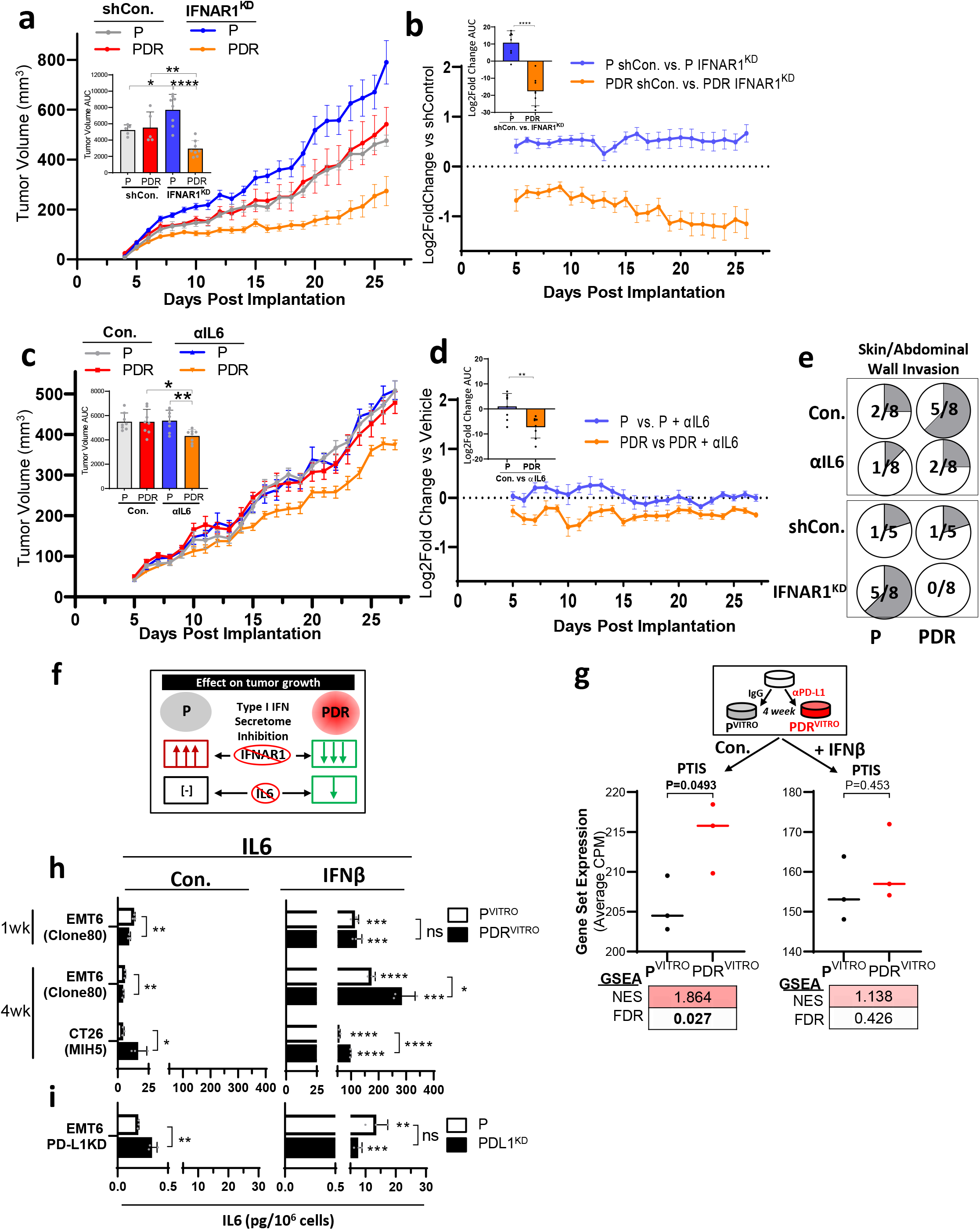
PDR tumor growth suppression by PTIS inhibition is partially PD-L1 regulated. **(a)** Orthotopic tumor growth of EMT6-P and EMT6–PDR, and respective IFNAR1^KD^ cell variants (n=5-8; Balb/c) with summary of AUC analysis (left-inset). **(b)** Log_2_ Fold Change analysis of data shown in (a) comparing IFNAR1^KD^ to respective controls. AUC analysis of comparisons shown (left-inset). **(c)** Orthotopic tumor growth of EMT6-P and EMT6–PDR treated with αIL6 antibody (n=5-8; Balb/c) with summary of AUC analysis (left-inset). **(d)** Log_2_ Fold Change analysis of data shown in (c) comparing αIL6 antibody treatment to respective controls. AUC analysis of comparisons shown (left-inset). **(e)** Metastasis and invasion of mice bearing EMT6-P and -PDR tumors (shown in a-b) with invasion in the peritoneum wall after PBS, αIL6, Con., or IFNAR1^KD^ (n=5-8). **(f)** Schematic summarizing the effect of type I IFN secretome inhibition in P and PDR tumors **(g)** Schematic showing generation of PDR^VITRO^ cell variants following αPD-L1 treatment *in vitro* for >4 weeks (top). Average CPM expression and GSEA of PTIS in EMT6 PDR^VITRO^ and PDR^VITRO^ following IFNβ stimulation RNAseq datasets (bottom). **(h-i)** IL6 expression after IFNβ stimulation in CM of **(g)** EMT6 and CT26 PDR^VITRO^ cell variants derived after 1 or >4 weeks of αPD-L1 (clone 80 or MIH5) treatment and **(h)** PD-L1^KD^ cell variants and P control cell variants (ELISA). Parental (P); PD-L1 Drug Resistant (PDR); IFNAR1 knockdown (IFNAR1^KD^); shRNA Vector Control (shCon); Area under the curve (AUC) in vitro derived Parental (R^VTTRO^); in vitro derived PD-L1 Drug Resistant (PDR^VITRO^); Gene Set Enrichment Analysis (GSEA); αPD-L1 Treatment-Induced Secretome (PTIS); normalized enrichment scores (NES); false discovery rate (FDR); Conditioned media (CM); PD-L1 knockdown (PDL1-KD). Cells were treated with 10ng/ml of IFNs and collected after 5 days for IL6 protein expression quantifications. * p<0.05, ** p<0.01, *** p<0.001, **** p<0.0001 except for GSEA where bolded numbers and (*) indicate FDR<0.25 αIL6 was administered at 100μg/mouse/3 days continuously, PBS was used as control (Con.). Primary tumor burden was assessed by caliper measurement. Quantitative data shown as mean ± SEM.

### Intracellular PD-L1 signaling partially regulates PTIS expression

Since PD-L1 has intrinsic cell signaling functions that can be regulated by IFN signaling^20,48^, it is possible that PTIS in tumor cells after acquired resistance may stem directly from PD-L1 inhibition. To test this, we targeted PD-L1 therapeutically and genetically *ex vivo* and assessed PTIS expression. First, EMT6 cells were treated with αPD-L1 (clone 80) or IgG *in vitro* for >4 weeks generating EMT6-P^VITRO^ and EMT6-PDR^VITRO^ cell variants **(Fig 5g; schematic shown)**. Transcriptomic analysis showed significant positive enrichment of the PTIS in EMT6-PDR^VITRO^ cells via GSEA **(Fig 5g; left panel);** with enrichment also observed after IFNβ stimulation, though this did not reach significance **(Fig 5g; right panel)**. Unlike *in vivo*-derived PDR cells, PTIS/ISGs in EMT6 PDR^VITRO^ cells were not consistently elevated at baseline, and IFNβ stimulation elevated some (but not all) PTIS components **(Fig S6a; Fig S6b shows heatmap comparison)**. When PD-L1 expression was knocked down in EMT6 cells (EMT6-PDL1^KD^; **Fig S6c**), PTIS/ISGs were partially elevated at baseline in EMT6-PDL1^KD^ cells, and IFNβ stimulation did not enhance PTIS expression **(Fig S6d; Fig S6e shows heatmap)**. For IL6 specifically, >4 weeks of exposure to two different mouse αPD-L1 antibodies (clone 80 and MIH5) and two different cell lines (EMT6 and CT26) yielded elevations of IL6 expression after IFNβ stimulation similar to *in vivo*-derived PDR cell variants, but these elevations at baseline were not consistent **(Fig 5g)**. Notably duration of treatment over shorter periods (1 week) or after PDL1^KD^ did not produce consistent IL-6 changes, marking a contrast with PD-L1 treatment resistant cells derived in *in vivo* mouse systems **(Fig 5g)**. Together, these results reveal that intrinsic PD-L1 signaling can account for some of the IFN-regulated PTIS observed in *in vivo*-derived resistance models, but this is partial and not consistent, indicating that tumor:immune cell interactions play a key role in PTIS adaptations in treatment-sensitive tumors.

## Discussion

A subset of cancer patients who are initially responsive to PD-L1 inhibitors will develop acquired resistance^59,60^. Mechanisms to explain why immunologically ‘hot’ tumors turn ‘cold’ remain complex as the microenvironment can adapt to treatment by relying on alternative checkpoints, inducing permanent T cell exhaustion, and recruiting/expanding an array of immunosuppressive cells – amongst many other changes attributed to host cell populations (reviewed in ref ^59,61^). But there is increasing evidence that the tumor also adapts to PD-L1 blockade^62,63^. Here, we examined the consequences of prolonged PD-L1 inhibition *in vivo* on tumor cells and identified a unique secretory signature that was associated with acquired resistance, enriched for numerous ISGs, and tightly regulated by type I IFN signaling. These tumor intrinsic adaptations were found to protect tumor cells from immune mediated cytotoxicity *directly*, via decreased sensitivity to lymphocytic attack, and *indirectly*, via a suppression of T cell activation. Importantly, resistant tumors were found to be uniquely vulnerable to IFN signaling disruption which, when targeted *in vivo*, could partly reverse immune-protective effects and enhance tumor growth inhibition. Together, these findings suggest that a consequence of chronic PD-L1 blockade includes a tumor secretory signature that may serve as a biomarker and molecular driver of acquired resistance in patients **(Fig 6)**.

**Figure 6:**
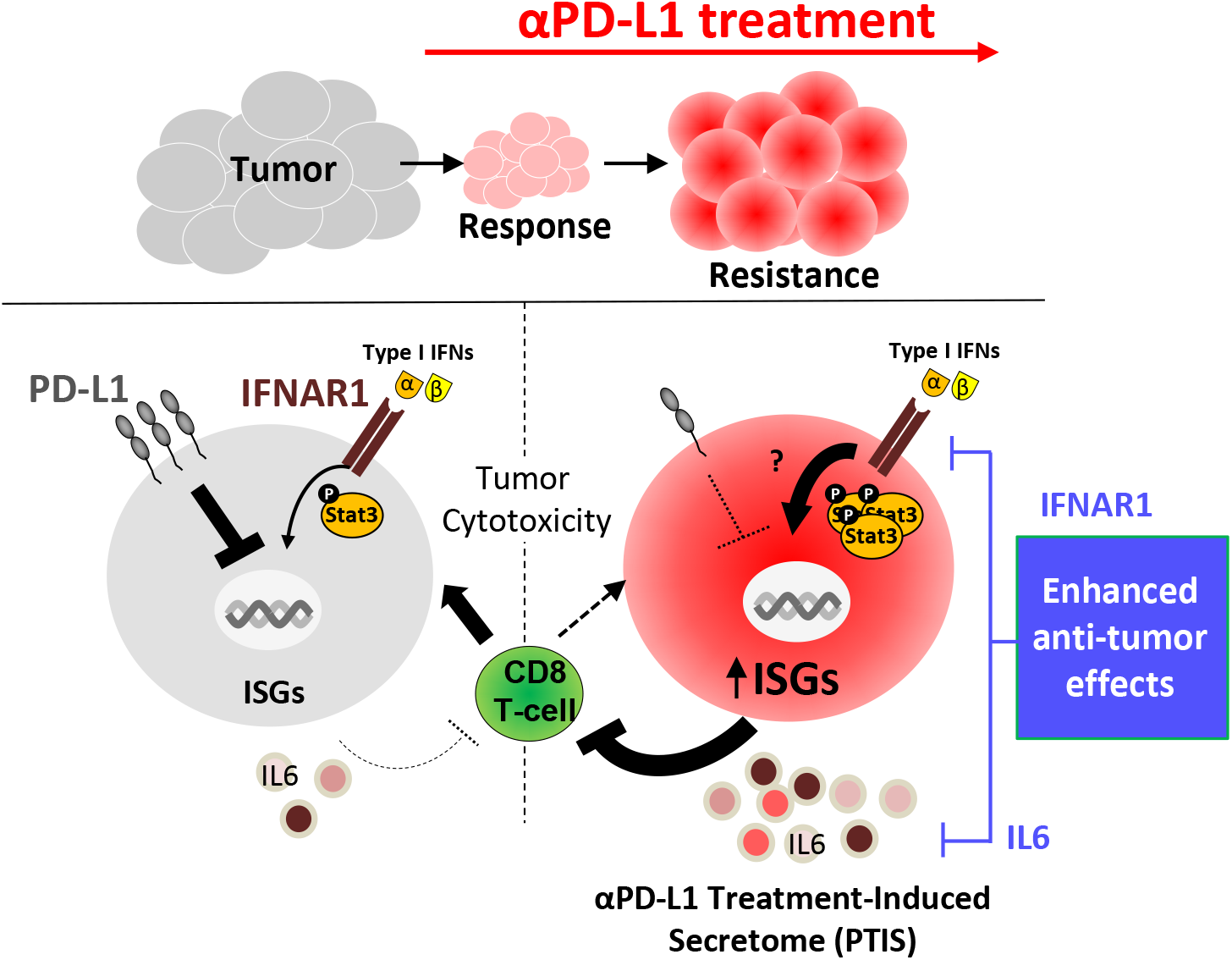
Proposed model of IFN-signaling ‘rewired’ tumor cells following acquired resistance to PD-L1 inhibition.

Currently the study of secretory changes in response to cancer treatments represent a phenomenon with broad implications for assessing (and improving) drug efficacy in patients^8,64^. Measuring systemic protein changes may provide clues into optimal drug dosing and give insight into emerging biological changes such as resistance^65,66^. For ICIs, levels of secretory proteins may correlate with different outcomes depending on disease site and duration of treatment. For instance, secreted circulating factors such as IL6, CXCL9, and CXCL10 have been found to initially increase in patients after PD-1 pathway inhibition and correlate with tumor stabilization/shrinkage, as measured by objective response rates (ORR) ^6,67–69^. But few studies have measured cytokine changes in patients after prolonged (long-term) treatment durations and assessed progression-free and overall survival (PFS/OS) outcomes^7,70^. Even when these are tested, results are often mixed. As an example, pre-treatment serum IL6 can correlate with improved initial response to nivolumab in a phase II trial for advanced melanoma (measured by ORR)^67^, but IL6 can also predict for worse long-term outcome to PD-1/L1 inhibitors in NSCLC (measured by PFS)^71^ and to atezolizumab/bevacizumab in hepatocellular carcinoma (measured by PFS and OS) ^72^.

Nevertheless, our studies suggest that PTIS signatures linked to acquired resistance and IFN signaling may warrant further investigation as a biomarker. Already transcriptomic analysis comparing clinical tumor tissues has shown that ISGs can increase in both tumor and non-tumor cell populations before and after ICI treatment in patients^16,33,36,73^. Our study showed that PTIS was enriched in 7 clinical and preclinical datasets involving αPD-L1 treatment-sensitive cancers, suggesting that testing PTIS in post-treatment biopsies may have utility as a predictor for level of tumor sensitivity. Yet challenges remain in using secretory changes a clinical biomarker in general. First, there is the difficulty in obtaining post-treatment biopsy tissue where treatment dose/duration are closely monitored, as clinical samples may be obtained long after treatment has ceased. Our study utilized published clinical data from αPD-L1 treated NSCLC^37^ and MCC^38^ tumors, but such data is rare. Perhaps the ideal setting to test secretory biomarkers would be in tissues/blood obtained after neoadjuvant PD-L1 inhibition, of which several trials are currently underway including in cutaneous melanoma patients treated with atezolizumab for 6 weeks (NCT04020809). In such testing, it is important to consider how genomic and proteomic data can be optimally paired to maximize potential predictive utility as gene expression of secretory proteins does not always translate to similar changes in protein expression, and vice versa^8,74^. For this reason, measuring circulating plasma protein measurements before and after anti-PD-L1 treatment in patients using multiplex immunoassay and proximity extension assays as a standardized assessment across studies may be considered to provide more consistent results^75–77^.

While biomarker testing may hold predictive information about treatment, one question raised by our study is whether PTIS factors themselves may represent a biological target to overcome resistance. Interestingly, the role of several PTIS proteins may largely be contextdependent as several have been linked to tumor-promoting *and* tumor-inhibiting immune responses. An example includes CXCL9 and CXCL10, which we found to be increased in PDR cells and, thus far, have both been primarily associated with anti-tumor immune responses, including trafficking of cytotoxic T cells^78–80^. But in different disease types and tumor models, these cytokines are also associated with recruitment of immune suppressive/tumor promoting T-regulatory cells^40–42^. In this regard, it is of particular interest that the PTIS was found to be enriched in non-tumor populations such as macrophages and T cells of Merkel cell carcinoma patients treated with avelumab **(Fig 2E)** - suggesting PD-L1 inhibition can likely induce ‘off-target’ host effects^8^. This may portend to the PTIS playing a role in immune related adverse events (irAEs) known to be induced by various ICI treatments and associated with the systemic increase in cytokines, such as IL6, IL1-RA, and CXCL10^81^. Indeed, PD-1 targeting agents have been known to induce cytokine release syndrome, which is an adverse effect characterized by fever, myalgias, malaise, and high levels of cytokines including IL6 and IFNs^82,83^. Assessment of the PTIS as a potential cause or consequence of these processes requires further investigation.

In this regard, assessing the role of IFN signaling on tumor growth and overall efficacy of ICIs remains complex. This is because IFNs can have opposing, and often contradictory, effects depending on several factors that include the type of cell, the duration of IFN exposure, and the stage of tumor progression ^17,84^. These stimulating/inhibiting effects IFNs can, in turn, influence ICI treatment response. For *inhibitory* effects, IFNs (mostly IFNγ) have historically been characterized to be integral for anti-tumor immunity by driving antigen presentation^37^ and chemokine secretion^85^ that are typically part of the immune-editing process in normal physiological conditions. IFNs can improve ICI responses as loss-of-function mutations in IFN signaling components (i.e. JAK1/2) have been identified in melanoma patients after pembrolizumab treatment relapse ^86^, and knockout of IFN pathway mediators such as Jak1, Stat1, Ifngr1 in tumors can weaken ICI treatment efficacy^87,88^. For *stimulating* effects, ICIs can induce an enhanced expression of tumor cell ISGs transcribing additional T-cell co-inhibitory ligands (e.g. TNFRSF14, LGALS9) where blockade of type I and II IFN signaling could reverse this effect and improve ICI responses^9^. Other tumor ISGs regulated by type I IFNs such as NOS2^14^ and CD38^16^ can also have immune-suppressive effects and have been implicated to promote resistance to PD-1 pathway blockade. In this context, one question is whether negative consequences of IFNs can be specifically targeted, without negating the positive IFN signaling effects. In this regard, Benci and colleagues proposed that ISGs promoting ICI resistance may be more associated with tumor cell expression^10^ and after chronic exposure to IFNs^9^, compared to the largely positive effects of IFNs on immune cells and after acute IFN exposure. Indeed, in murine tumor models, success of ICI treatment depends on a ‘display-on/fast-off’ kinetic of IFNβ signaling where fast reduction of ISG expression is associated with responding tumors^89^. Therapeutically, sequential treatment of an ICI followed by a JAK inhibitor was found to sensitize IFN-driven resistant tumors to ICI treatment^9^. Such treatment strategies are now being tested in clinical trials (NCT03425006).

This raises the question of whether disruption of the PTIS in PDR cells should involve targeting the secretory-controlling IFN-signaling machinery or targeting ISG secretory products directly, thereby potentially avoiding disruption of IFNs anti-tumor functions. In our study, both approaches were evaluated. In the first, IFNAR1 knockdown in PDR cells was found to effectively reverse IFNβ-mediated PTIS factor increases and reverse direct (and indirect) immune suppressive effects in co-culture studies with mouse splenocytes. Notably these studies were conducted using CD3/CD28 agonistic antibodies to bypass T cell activation, suggesting that PDR secretomes were capable of suppressing T cell activation independent of antigen presentation. This may explain why PDR-IFNAR1^KD^ tumors were found to grow slower than PDR tumors *in vivo*, suggesting that constant PD-L1 blockade confers a unique immune-promoting vulnerability in tumor cells, despite the intact antigen presentation machinery that we found provides an immune-suppressive counterbalance ^37^. However, our results also show that IFNAR1^KD^ tumors grew much faster than parental controls, emphasizing the sometimes contradictory role of IFNAR signaling in cancer controlling immunosurvellance^17^ which might be exacerbated by anti-PD-L1 treatment and, as a consequence, may introduce potential challenges of intracellular IFN signaling inhibition strategies.

For targeting an individual PTIS factor, we evaluated IL6 - which is a specific component of the PTIS found to be consistently increased in PDR cells and then further enhanced after IFNβ stimulation. IL6 is known to activate a multitude of tumor promoting effects that include (i) enhancing expression of pro-angiogenic factors in tumors cells (e.g. VEGF, IL1β, IL8^90^), (ii) suppressing antigen presentation from dendritic cells ^91^, (iii) promoting pro-tumorigenic macrophage phenotypes^92^, and (iv) suppressing anti-tumor functions of CD4+ T cells ^93^ (amongst many others^94^). Trials are currently underway testing IL6 and PD-1 pathway inhibitor combinations for improved anti-tumor efficacy and irAEs (NCT03999749, NCT04258150)^95,96^. Indeed, our results showing that IL6 inhibition can lead to enhanced growth suppression in the acquired resistance setting adds to the growing literature supporting anti-IL6/PD-L1 combination/sequencing strategies. However, it should be noted that these effects were largely modest, suggesting targeting multiple PTIS factors simultaneously may yield more robust outcomes. In favor of this, preclinical models involving inhibition of CCL2^97^, CCL5^98^, NOS2^14^, and SERPINB9^15^ pathways have all shown benefits when combined with various ICIs and may be explored to improve IL6 inhibitory strategies after PD-L1 treatment failure.

A unique feature of our studies is the use of *in vivo*-derived models to evaluate acquired resistance, which served to mimic the evolution of treatment-sensitive tumor progression to eventual insensitivity. For ICIs, acquired resistance is important but surprisingly difficult to recapitulate in preclinical systems. Murine tumor models have translational value by providing an accurate assessment of tumor-immune interactions, yet the majority do not demonstrate an initial response to single agent ICI treatment (previously reviewed^99,100^). This is perhaps best demonstrated by Mosely et al., where six of the most commonly used syngeneic tumor models showed that only two (CT26 and RENCA) were sensitive to anti-CTLA-4 therapy and only one (CT26) was sensitive to anti-PD-L1^34^; with later studies suggesting that CT26 response to PD-L1 inhibition may depend on antibody clonality^101^ and duration/timing of treatment schedules^102^. Explanation for this limited efficacy is still unclear and may be model-dependent due to variations on T cell infiltration^103^, infiltration of immunosuppressive cell types^34^, tumor mutation burden^34^, and both tumor and non-tumor host expression of immune checkpoint molecules^104^. Our studies identify EMT6 as a treatment-sensitive model for acquired resistance that can be used to confirm molecular changes with clinical datasets to establish translational relevance. As ICIs are approved in new disease indications, the use of such models will be increasingly important for studies examining mechanisms, biomarkers, and therapeutic solutions for ICI efficacy.

Lastly, results from our studies suggest tumor-intrinsic signaling can lead to extrinsic secretory changes after PD-L1 inhibition, potentially stemming from changes caused by PD-L1 directly. Though often overlooked, PD-L1 has several tumor intrinsic functions that have recently been identified to regulate mTOR/AKT^105,106^, MAPK^107^, STAT3/Caspase 7^20^, integrin β4^108^, MerTK^109^, and BIM/BIK^110^ signaling, among others. Recently, Gato-Canas and colleagues reported that conserved motifs of PD-L1 cytoplasmic domains (RMLDVEKC and DTSSK) block STAT3/Caspase 7 cleavage and, in turn, can control type I IFN-induced cytotoxicity^20^. This result introduced PD-L1 as a ‘molecular shield’ that can protect tumor cells from T cell cytotoxicity caused by various treatments^111^, including IFNs^20,112^. Our results suggest that some, but not all, of the IFN-enhanced PTIS observed after PDL1 resistance may be caused by intrinsic PD-L1 signaling. Genetic disruption of PD-L1 led to an enrichment of several PTIS factors but, unlike *in vivo*-derived PDR cells, IFNβ stimulation did not lead to a further increase. For *in vitro*-derived PDR cells generated by chronic exposure to PD-L1 antibody, the opposite finding occurred, indicating that the molecular ‘rewiring’ of tumor cells after acquired resistance to PD-L1 requires the complex molecular and cellular changes from host processes that indirectly alter tumor cell PD-L1 and IFN signaling. For instance, PD-L1 inhibition is known to modulate functions of immune cells such as T-, NK-, and myeloid cell populations, amongst several others to drastically remodel the tumor microenvironment^113–115^ as well as modulate local and systemic cytokine and growth factor changes that include IFNs^8^. Overall, our results align with previous studies suggesting that PD-L1 has both activating and inhibitory domains on tumor cell IFN signaling that may be impacted differently by genetic or therapeutic strategies to block signaling^20^. Nevertheless, our findings also highlight the challenges of studying acquired resistance mechanisms to PD-L1 inhibition and the critical need for appropriate *in vivo* systems to bridge findings from *in vitro* studies involving PD-L1 functional signaling, particularly as it pertains to translational significance of therapeutic relapse as it occurs in patients.

Taken together, our results show that tumor cells following acquired resistance to PD-L1 blockade can express an ISG enriched secretory profile associated with diminished sensitivity to immune cell cytotoxicity. Therapeutic approaches involving inhibition of PTIS components or IFN regulators may have enhanced benefit after resistance to PD-L1 inhibition.

## Acknowledgements

We would like to thank K. Eng for consultations and guidance on bioinformatic analysis of singlecell sequencing data; and A. Tracz for animal work assistance. Select cell lines used in this study were kind gifts from various laboratories. These include EMT6 (A. Gukov) and B16 (D. Escors). We thank M. Azuma for providing the MIH5 hybridoma for antibody production (see Methods) and M. Oberst at AstraZeneca for providing the Clone 80 antibody.

## Author Contributions

Conceptualization, YS, JMLE; Methodology, YS, MD, MM, ML, SB, JMLE; Investigation, YS, MD, MM, JWH, AD, IP, SA, JMLE; Formal Analysis, YS, MM, MD, JMLE; Visualization, YS, MM, JMLE; Supervision, JMLE; Funding Acquisition, YS, JMLE; Writing – Original Draft, YS, JMLE; Writing – Review and Editing, YS, MD, MM, JMLE.

## Materials and Methods

### CONTACT FOR REAGENT AND RESOURCE SHARING

Further information and requests for resources and reagents should be directed to and will be fulfilled by Lead Contact, John M.L. Ebos.

#### Cell lines

Cell used in this study include: Mouse mammary carcinoma EMT6 (from A. Gudkov, Roswell Park Comprehensive Cancer Center, RPCCC), colorectal carcinoma CT26 (A. Gudkov), and mouse kidney RENCA (from R. Pili, RPCCC). Cells were maintained in RPMI (Corning cellgro #10-040-CV) supplemented with 5% v/v FBS (Corning cellgro; 35-010-CV). All cells were maintained at 37°C with 5% CO2 in a humidified incubator.

#### Drug and recombinant protein concentrations

IgG1 (NIP228, AstraZeneca), IgG2a (I-1177, Leinco Technologies Inc), αPD-L1 (clone 80, AstraZeneca), αPD-L1 (MIH5, from M. Azuma, Tokyo Medical and Dental University^116^) and anti-IL6 (BE0046/MP5-20F3, BioXCell) were prepared as follows: For *in vivo* experiments: αPD-L1 (Clone80) and anti-IL6 (MP5-20F3) were diluted in PBS and administered by intraperitoneal injection at (250μg/mouse/3days) or (100μg/mouse/3days) respectively. Tumor-related differences between any vehicle or IgG groups were not observed. I*n vitro*, IgG (NIP228 or I-1177) and αPD-L1 (clone80 or MIH5) in PBS was directly added to media for maintenance at a concentration of 0.5μg/ml; anti-IL6 was used at a concentration of 10μg/ml; recombinant IFN-alpha-2 (50525-MNAY, Sino Biological), IFN-beta (50708-MCCH, Sino Biological), IFNγ (315-05, Peprotech) were used at 10ng/ml.

#### shRNA knockdown studies

For production of IFNAR1 and PD-L1 knockdown lentivirus, pLKO.1-puro shRNA plasmid DNA was isolated from bacteria glycerol stocks (IFNAR1: TRCN0000301483, PD-L1: TRCN0000068001; Sigma Aldrich) using E.Z.N.A.^®^ Plasmid Mini Kit I (Omega Bio-tek, Inc.). To produce lentiviral media, 293T cells were transiently co-transfected with DNA from the lentiviral pLKO.1-puro shRNA plasmid and psPAX2 and pMD2.G packaging plasmids using LipoD293™ Transfection Reagent (SignaGen Laboratories.) Conditioned media containing virions was harvested after 24 and 48 hours, filtered through a 0.45-μm membrane, and used to infect EMT6-P or PDR cells. Cells were infected with the shRNA and vector controls by spin inoculation at 600 × g for 45 min at room temperature in the presence of 5ug/ml polybrene. Viruses were removed after an additional 6hr incubation at 37°c/5% CO2 and cell culture media was replaced. Puromycin selection was then conducted for 2 weeks at 2μg/ml until stably infected cells were generated. Knockdown was confirmed via flow cytometry analysis.

#### Mouse tumor models

##### Study Approval

Animal tumor model studies were performed in strict accordance with the recommendations in the Guide for Care and Use of Laboratory Animals of the National Institutes of Health and according to guidelines of the IACUC at RPCCC (Protocol: 1227M).

##### Orthotopic Tumor Implantations

EMT6 (5×10^5^ cells in 100μl RPMI), RENCA (4×10^4^ cells in 5μl 1:1 RPMI:Matrigel) were implanted orthotopically into the right inguinal mammary fad pad or left kidney subcapsular space respectively in 6-8 week old female Balb/c mice. Isoflurance (anesthesia) and buprenorphrine (analgesic) were used during all surgical implantations. Mammary fat pad tumors were measured using Vernier calipers and volumes were calculated using the formula (width^2^ x length) x 0.5. Kidney luciferase expressing tumors were assessed for bioluminescence activity bi-weekly. All animals were assessed 2-3 times daily by veterinary staff or personnel approved by IACUC for pre-defined endpoints. Institutional endpoints included primary tumor-based morbidities (>2000mm^3^ volume) and metastasis related morbidities (labored breathing, 20% weight loss, cachexia, limb paralysis). All mice were randomized before implantation.

#### Resistance cell derivation and maintenance

For *in vivo*-derived PDR cell variants, mice were orthotopically implanted with EMT6 or RENCA and treated with αPD-L1 (Clone80) until institutional endpoint. For both EMT6 and RENCA, parental (P) cell lines were obtained from IgG-treated mice and used as controls. All variants were selected from primary tumors which were minced, enzymatically digested (Miltenyi Biotics; 130-095-929), and then placed in RPMI media (supplemented with 5% v/v FBS, 100IU/ml penicillin and 1000 μg/ml streptomycin) with IgG (NIP288) for P variants or with αPD-L1 (clone80) (0.5μg/ml) for PDR variants. Antibiotics were then removed 1 week after in vivo cell selection. For derivation of PDR^VITRO^ cell variants, EMT6 and CT26 cells were treated with αPD-L1 antibodies (clone 80 at 0.5μg/ml or MIH5 at 0.5μg/ml, respectively) for >4 weeks.

#### Cell proliferation assay

Proliferation was examined using CellTiter 96 Aqueous Non-Radioactive cell proliferation (MTS) assay (Promega; G1112). For 5 day growth studies, 200 cells/well were plated in 48-well plates. The next day, cells were treated with recombinant IFNs or anti-IL6. Treatments were replaced every 2 days or removed daily for MTS measure of viability. RPMI +5% FBS was mixed with MTS per manufacturer instructions and added to cells at timepoints. After 2 hour incubation, optical density was measured at a wavelength of 490nm (Bio-Rad xMark).

#### RNA isolation

Cells were plated at 80,000 cells/well in a 6 well plate with corresponding treatments as indicated. 48 hours later, total RNA was isolated using QIAshredder (QIAGEN; 79654) and RNase mini kit (QIAGEN; 74104). Genomic DNA was then digested using DNaseI (QIAGEN; 79254) per manufactuer instructions. RNA concentration was determined using nanodrop 2000c (Thermo Scientific) before RNAseq and PCR analysis.

#### qRT-PCR

For reverse transcription using iScript cDNA synthesis kit (Bio-rad; 170-8891),1μg RNA was used according to the manufacturer’s instructions. qRT-PCR was performed using iTaq SYBR Green Supermix (Bio-rad; 1725121). Thermocycling parameters were: 10 min at 95oC, 15 sec at 95oC, 40 cycles at 95oC for 15 sec at 95oC and 1 min at 60oC, 1min at 95oC, followed by a melting curve: 55 to 95oC with increments of 0.5oC for 5 sec. Relative gene expression was calculated using the formula 2^[-CT(House Keeping Gene)-Ct(Gene of Interest)]^, with CT representing the fixed threshold cycle value for fluorescent signal. Gapdh and Actb were used for housekeeping genes. Oligonucleotides were purchased from Integrated DNA Technologies (IDT) **(Table S3).**

#### Proteome profiler (cytokine) array

Cells were lysed with lysis buffer 17 (R&D Systems; 895943) supplemented with protease cocktail (Fisher Scientific, PI78430). Total protein levels were quantified with DC protein assay (Bio-Rad; 500-0112). 200μg of total mouse protein samples were analyzed respectively with a Mouse XL Cytokine Array Kit (R&D Systems; ARY028) per manufacturer instructions. Membranes were exposed to X-ray films, which were imaged (digitized) with ChemiDoc System (Bio-Rad) and analyzed with Image Lab Software (Bio-Rad).

#### ELISA analysis

Cells were lysed with lysis buffer I (20mM Tris (pH7.5), 127mM NaCl, 10% Glycerol, 1% v/v NP40 (Igepal), 100mM NaF, 1mM Na3VO4) and protein concentrations were quantified with DC protein assay. For conditioned media collection, cells were counted for normalization. IL6, phospho-STAT3, and PD-L1 were measured using mouse IL6 ELISA (431304, Biolegend), mouse phospho-STAT3 ELISA (7300C), and mouse PD-L1 Duoset ELISA (DY1019-05).

#### Western blot analysis

Cells were lysed with lysis buffer II (50mM Tris (pH8), 2% w/v SDS, 5mM EDTA, 3mM EGTA, 25mM NaF, 1mM Na_3_VO_4_) supplemented with Halt™ protease inhibitor (Thermo Fischer Scientific 78429). Lysates were sonicated for 2 seconds and total protein concentration was quantified with DC protein assay. Proteins samples were prepared with 1/5 volume of 5x SDS-PAGE sample buffer (250mM Tris pH6.8, 10% w/v SDS, 25% v/v glycerol, 500mM DTT, and bromophenol blue). Proteins (40 μg per lane) were resolved by SDS-PAGE, electrotransferred to Immobilon-P membrane, and incubated with a primary antibody diluted as recommended by the manufacturer. Membranes were then probed with a horseradish peroxidase-conjugated secondary antibody (Promega W4011 and W4021) and protein signals were developed using the Pierce ECL Western blotting substrate (Thermo Scientific; 32106). X-ray films were imaged (digitized) with ChemiDoc System and analyzed with Image Lab Software. Primary antibodies were purchased from Cell signaling (phospho-STAT1 Tyr701 9167S, STAT1 14994, phospho-STAT3 Tyr705 9145T, STAT3 9139T) and Sigma Aldrich (β-actin, A5441).

#### Splenocyte Division and Activation Assays

Spleens were harvested from Balb/c mice, mechanically dissociated by passing through a 70μm filter, and collected in complete RPMI media (supplemented with 10% heat-inactivated FBS, 1% non-essential amino acids, 1% sodium pyruvate, 1% penicillin/streptomycin, and 0.1%β-mercaptoethanol). Splenocytes were then treated with RBC lysis buffer (Biolegend, 420301) and incubated overnight. The next day, splenocytes were stained with CFSE (Biolegend, 423801), stimulated with anti-mouse CD3 (Biolegend, 100202) and CD28 (Biolegend, 102102) according to manufacturer recommendations. To generate conditioned media, 7 x 10^6^ tumor cells were plated with 10ml of RPMI (supplemented with 5% FBS) in a 10-cm dish. After 72 hours, conditioned media was collected and passed through a 0.2μm filter. Splenocytes were then plated in a 1:1 ratio of complete RPMI:conditioned media at 400,000 cells per well in a 96-well plate. After 72 hours of incubation splenocytes were treated for 5 hours with activation cocktail with brefeldin A1 (Biolegend, 423303) before staining for CD45 (Biolegend, 103128), CD8b.2 (Biolegend, 140416), Granzyme B (Biolegend, 515408), IFNγ (Biolegend, 505808), and CD69 (Biolegend, 104514) according to manufacture instructions.

For tumor cell co-culture experiments, 1×10^4^ tumor cells were plated in a 24-well plate per well and allowed to adhere overnight. The next day, 1×10^6^ stained and stimulated splenocytes were added per well. After 72 hours of incubation, splenocytes were processed as described above and adherent tumor cells were concurrently collected for Annexin V (Biolegend, 640920) and 7-AAD staining (Biolegend, 420404) via flow cytometry according to manufacturer instructions.

#### Flow cytometry analysis of cell surface proteins

Cells were plated at 80,000 cells/well in a 6 well plate. After two days, cells were collected by accutase (Biolegend, 423201) and analyzed by flow cytometry for H-2Kb/H-2Db/H-2Dd (Biolegend, 114613) expression according to manufacture instructions.

#### Whole transcriptome expression analysis

RNA sequencing for tumor tissue-derived EMT6-PDR, and tumor cell line-derived EMT6-PDR^VITRO^ and RENCA-PDR cells, were performed utilizing the Genomic shared resource at RPCCC as previously described ^117^. Sequencing library were prepared with TruSeq Stranded mRNA kit (Illumina Inc), from 1μg total RNA, according to manufacturer’s instructions. PolyA selection, RNA purification, fragmentation and priming for cDNA synthesis was performed. Using random primers, fragmented RNA was then reverse transcribed into first-strand cDNA. RNA template was then removed, a replacement strand was synthesized and dUTP was incorporated in place of dTTP to generate ds cDNA. ds cDNA was separated from second-strand reaction mix using AMPure XP beads (Beckman Coulter) resulting in blunt-ended cDNA. One ‘A’ nucleotide was added to the 3’ ends of the blunt fragments. Multiple indexing adapters, containing one ‘T’ nucleotide on the 3’ end of the adapter, were ligated to the ends of the ds cDNA, preparing them for hybridization onto a flow cell. Adapter ligated libraries were amplified by PCR, purified using Ampure XP beads, and validated for appropriate size on a 4200 TapeStation D1000 Screentape (Agilent Technologies, Inc.). The DNA libraries were quantitated using KAPA Biosystems qPCR kit, and were pooled together in an equimolar fashion, following experimental design criteria. DNA library pool was denatured and diluted to 2.4pM with 1% PhiX control library added. The resulting pool was then loaded into the appropriate NextSeq Reagent cartridge, as determined by the number of sequencing cycles desired, and sequenced on a NextSeq500 following the manufacturer’s recommended protocol (Illumina Inc.). Sequencing quality control was assessed using FASTQC v0.11.5 (http://www.bioinformatics.babraham.ac.uk/projects/fastqc/). Reads were aligned to the mouse genome GRCM38 M16 (genocode) using STAR v2.6.0a^118^ and post-alignment quality control was assessed using RSeQC v2.6.5^119^. Aligned reads were quantified using RSEM v1.3.1 ^120^. Counts from RSEM were then filtered and then upper quartile normalized using R package edgeR. Data from RENCA and EMT6 studies will be deposited in GEO prior to manuscript publication.

#### Gene Ontology Analysis/Cytoscape

Differentially expressed genes with products located in extracellular regions were identified using gene ontology databases (GO:00005576) as previously described^8,74^. GO Biological processes terms were then assessed using ClueGo via Cytoscape v3.7.2 and significantly enriched terms and corresponding Kappa scores were plotted based on p-values.

#### Gene Set Enrichment Analysis

Gene set enrichment analysis (GSEA) was conducted to assess comparisons for molecular pathways and gene set correlations. A rank list was first generated using log2(fold change) gene expression data obtained from limma anlaysis. GSEA-Preranked was then conducted using a gene-set permutation type with 1000 random permutations to obtain normalized enrichment scores (NES) and false discovery rate (FDR) q-values.

#### Interferome Analysis

Genes of interest were assessed in the Interferome Database (http://www.interferome.org/interferome/home.jspx) ^28^ to examine for evidence of regulation by IFN signaling. Parameters interferome type, subtype, treatment concentration, treatment time, *in vivo/in vitro*, species, system, organ, cell, cell line, normal/abnormal were set to “any”, fold-change thresholds were set to 1.5.

#### Identification of a PD-L1 treatment-induced secretory (PTIS) gene signature

To identify gene signatures associated with secretory products unique to PDR tumor cells, we created a preliminary list of PTIS genes (PTIS ‘preliminary’) by extracting secretory genes (**Table S2**). Using this preliminary list, an enriched PTIS gene signature was created from comparisons of EMT6-P and EMT6-PDR using various transcriptomic and proteomic assays (**Table S2; Figs S1a-d**). These included individual assessment of differentially expressed secretory genes from bulk RNAseq transcriptomics data (**Fig S1**; with fold-change >2 or <-2) and proteomic comparisons of cell lysate using a proteome profile (cytokine) array (**Fig S1b**). The PTIS was further enriched with targets added based on individual comparisons of EMT6-P and EMT6-PDR variants using qRT-PCR (**Fig S1c**) and ELISA (**Fig S1d**) assays. The final enriched PTIS signature was based on genes validated via at least 2 methods and with consistent results (i.e., excluding factors increased in a gene assay, but decreased in a protein assay, or vice versa) (**Fig S1e**). For transcriptomic comparisons of P and PDR variants (EMT6) shown in **Fig S2**, a similar methodology was used to generate a PTIS gene signature comprised only of genes downregulated (PTIS^DOWN^; **Table S2 and Supplemental Results**).

#### CIBERSORT/ImmuCC Analysis

Cibersort tissue deconvolution was performed using the ImmuCC signature to obtain absolute score for various cell types^51,52,121^. From 25 immune cell type signatures, no values were detected for eosinophil cells, CD4 Memory T cells, Neutrophil cells, and Plasma Cells; and thus, were excluded for quantification and analysis.

#### Confirmation of PTIS signature enrichment in published datasets

Previously published clinical and preclinical datasets derived from studies after treatment or after acquired resistance to PD-L1 inhibition were obtained from the Gene Expression Omnibus (GEO, www.ncbi.nlm.nih.gov/geo/) or database of Genotypes and Phenotypes (dbGaP, https://www.ncbi.nlm.nih.gov/gap/).

##### Sceneay et al. 2019 (GSE130472)

In this study, whole tissue RNA-seq (Illumina NextSeq500 with paired-end 75bp reads) were performed on 4T1 orthotopically implanted mammary tumors in 8-12 week (young; responsive) or >12months (old, nonresponsive) Balb/c mice treated with αPD-L1 (clone 10F.9G2) or isotype (Clone LTF-2). Tumors were collected when caliper volumes were no larger than ~150mm^3^ after 3 doses of antibody treatment.

##### Lan et al. 2018 (GSE107801)

In this study,_whole tissue RNAseq was (Illumina Hiseq 2500) conducted on EMT6 orthotopically implanted mammary tumors in Balb/c mice treated with αPD-L1 or isotype control. Tumors were collected 6 days after 3 doses of treatment given daily starting 20 days after implantation.

##### Efremova et al. 2018 (GSE93017)

In this study, whole tissue RNAseq (Ion Torrent Proton) was conducted on MC38 subcutaneously implanted tumors in C57Bl/6 mice treated with αPD-L1 (Clone10F.9G2) or isotype (Clone LTF-2) every 3-4 days until day 14 after implantation.

##### Gettinger et al. 2018 (phs001464.v1.p1)

In this study, pre-treatment and post-treatment/acquired resistant biopsies were obtained from patients receiving various immune checkpoint inhibitor treatments (PD-L1, PD-1, CTLA-4 targeted therapies) for RNA-seq analysis (Illumina HiSeq2500) from formalin fix paraffin embedded samples. For validation analysis, all pretreatment samples were compared to samples after acquired resistance to PD-L1 inhibition. Note: the name of the PD-L1 inhibitor was not provided in this publication.

##### Paulson et al. 2018 (GSE117988, GSE118056)

In this study, single cell RNAseq analysis on tumor tissues were conducted on an untreated patient biopsy (Illumina HiSeq 2500) and a patient biopsy at acquired resistance (unmatched) after αPD-L1 (avelumab), MCPyV-specific T cells, and radiation (Illumina NovaSeq 6000). Processed data were obtained from the GEO database. R packages SingleCellExperiment^122^, scater^123^, limma^124^, and Rtsne were used for analysis. Counts were quartile normalized and converted to counts per million (CPM). Clustered enrichment analysis was then conducted using markers PTRC/CD45 (tumor cells), CD3D (T cells), and CD68 (Macrophages) similar to previously described^38^.

PTIS signature expression levels in these datasets were compared by average counts per million (CPM) levels or GSEA enrichment (defined by FDR ≤ 0.25).

#### Statistical analysis

Analysis was conducted using the GraphPad Prism software package v8.4.0 (GraphPad software Inc., San Diego, CA) and R v3.6.0 through RStudio v1.1.463(Integrated Development for R; RStudio, Inc., Boston, MA URL http://www.rstudio.com/). For *in vivo* studies, results are represented as mean ± standard deviation (SD) or standard error of mean (SEM), as indicated. Kaplan-Meier methods were utilized for analysis of percent to institutional endpoint curves. Fold change differences between treatment control groups were assessed via two-way ANOVA. For all results, comparisons between two groups were made with Student’s two-tailed unpaired t-test, whereas one-way ANOVA was used for comparison of more than two groups. Tumor volume and bioluminescence measurements were compared for specified time points. A minimum FDR value of 0.25 was use for GSEA analysis (as indicated as described by the user guide) and significance level of 0.05 was used for all other analyses.

