## Supplemental Material for "Acquired resistance to PD-L1 inhibition is associated with an enhanced type I IFN-stimulated secretory program in tumor cells"

### Supplementary Results:

A downregulated PTIS signature is variably expressed in clinical and preclinical models sensitive to αPD-L1 treatment

#### Supplementary Results

##### A downregulated PTIS signature is variably expressed in clinical and preclinical models sensitive to αPD-L1 treatment

Concurrent with the generation of a αPD-L1 treatment-induced secretome (PTIS) signature comprising of *up*regulated genes following acquired resistant (from Fig 2A, Fig S1), we also generated a signature comprised only of genes significantly *down*regulated (**Fig S2a; PTIS$^{\mathbf{DOWN}}$**). Similar to the PTIS, the PTIS$^{DOWN}$ was comprised primarily of IFN-regulated genes (3 out of 4 total). Using published preclinical datasets representing αPD-L1 treatment-*sensitive* and treatment-*insensitive* tumor models (described in Fig 2b), enrichment for the PTIS$^{DOWN}$ was found to be variable with no significant signature expression trends (measured by CPM) found and only 1 of 5 (*insensitive;* RENCA-PDR) was found to have significant negative GSEA enrichment (**Fig S2b and S2c**). Analysis of PTIS$^{DOWN}$ signature expression in clinical datasets showed decreases in NSCLC PDR samples **(Fig S2d; significance not reached)**; while clustered analysis of MCC avelumab treated samples showed variable, but significant changes in enriched tumor cell (decreased expression) and macrophage clusters (increased expression) (**Fig S2e)**. Together these results suggest the PTIS signature representing *up*regulated genes are more representative tumor-intrinsic molecular changes that occur *in vivo* after acquired resistance.

### Supplementary Figures:

Figure S1: Generation of an enriched PD-L1 treatment induced secretome (PTIS) signature.

Figure S2: Analysis of PTIS$^{DOWN}$ in published preclinical and clinical datasets after PD-L1 inhibitor treatment

Figure S3: *In vitro* proliferation of EMT6-P and PDR cells after IFN signaling stimulation

Figure S4: Extended Data from Fig 3. - Protein analysis of ISGs in PDR cells

Figure S5: Extended Data from Fig 4. - Splenocyte activation following incubation with PDR cells and conditioned media.

Figure S6: Extended data from Fig 5- PTIS expression in PDR$^{VITRO}$ and PD-L1$^{KD}$ models

**Figure S1**

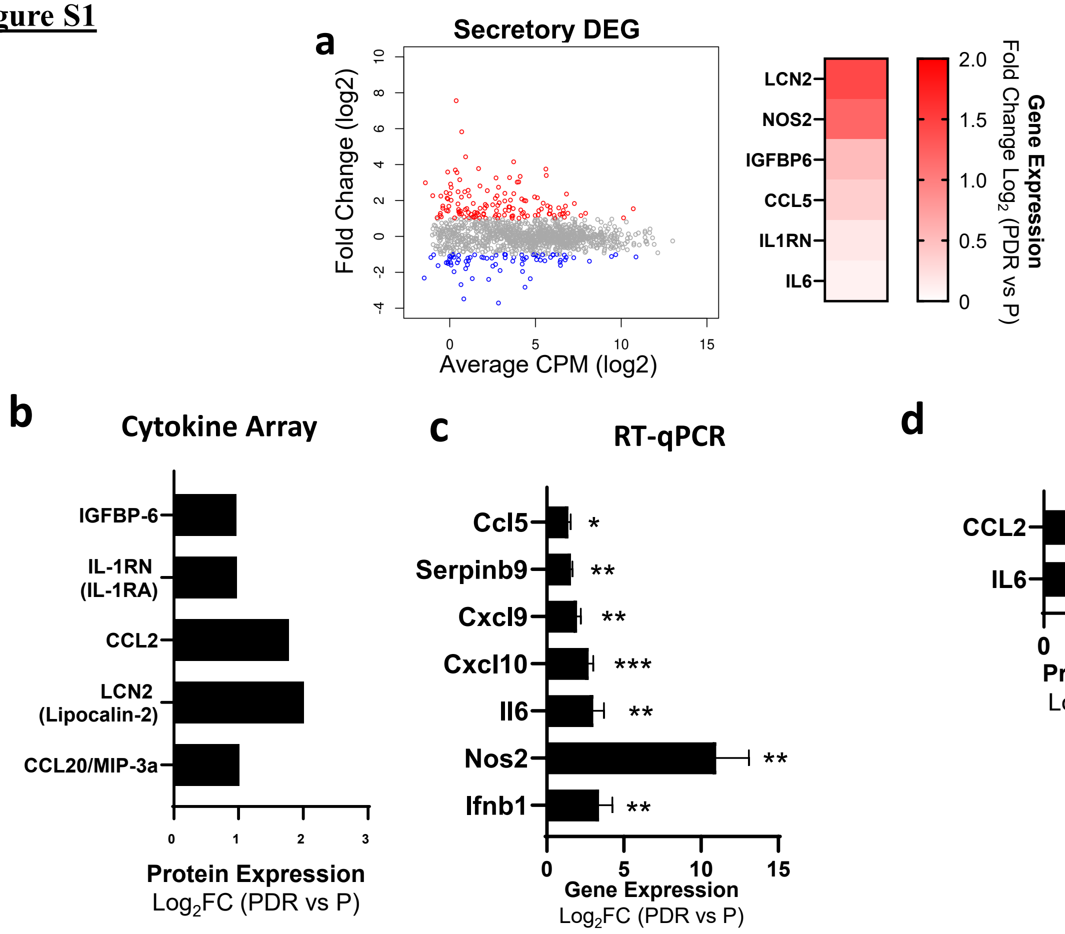


**e**

| PTIS | RNAseq | Protein Array | qRT-PCR | ELISA |
|---|---|---|---|---|
| CCL2 | | • | | • |
| CCL20 | | • | | |
| CCL5 | • | | • | |
| CXCL10 | | | • | |
| CXCL9 | | | • | |
| IFNB1 | | | • | |
| IGFBP6 | • | • | | |
| IL1RN | • | • | | |
| IL6 | • | | • | • |
| LCN2 | • | • | | |
| NOS2 | • | | • | |
| SERPINB9 | | | • | |

**Figure S1: Generation of an enriched PD-L1 treatment induced secretome (PTIS) signature.**

**(a)** Primary identification of differentially expressed secretory genes ($Log_2$ [FC] ≤ -2 or ≥ 2) in EMT6-PDR as summarized by dot plot (left) and heatmap (right) showing selected PTIS genes associated with IFN signaling. Red = upregulated; blue = downregulated. PTIS signature generation described in Table S2 and Methods.

**(b-d)** Secondary identification and validation of PTIS targets by cytokine array (b), RT-qPCR (c), and ELISA (d).

**(e)** IFN-enriched PTIS gene list used in GSEA screening of published data involving PD-L1 inhibitor treatment (shown in Fig 2a).

*Parental (P); PD-L1 Drug Resistant (PDR); Fold-Change (FC), Differentially expressed genes (DEG); PD-L1 Treatment Induced Secretome (PTIS)*

#### Figure S2

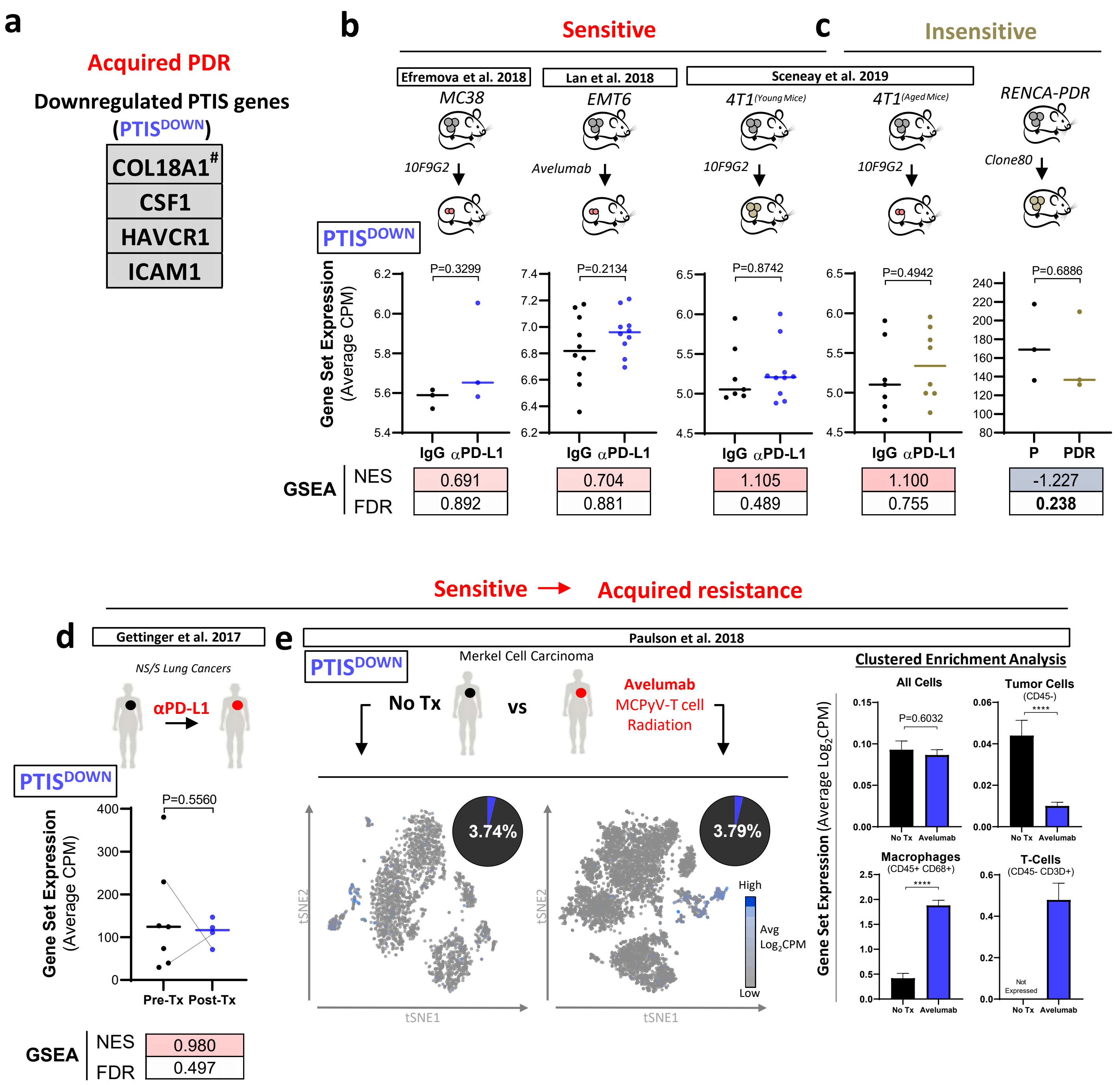

a
Acquired PDR
Downregulated PTIS genes
(PTIS DOWN)
COL18A1#
CSF1
HAVCR1
ICAM1
b
Sensitive
c
Insensitive
Efremova et al. 2018
Lan et al. 2018
Sceneay et al. 2019
MC38
EMT6
4T1 (Young Mice)
4T1 (Aged Mice)
RENCA-PDR
10F9G2
Avelumab
10F9G2
10F9G2
Clone80
PTIS DOWN
Gene Set Expression (Average CPM)
P=0.3299
P=0.2134
P=0.8742
P=0.4942
P=0.6886
IgG αPD-L1
IgG αPD-L1
IgG αPD-L1
IgG αPD-L1
P PDR
GSEA
NES
FDR
0.691
0.892
0.704
0.881
1.105
0.489
1.100
0.755
-1.227
0.238
Sensitive → Acquired resistance
d
Gettinger et al. 2017
NS/S Lung Cancers
αPD-L1
PTIS DOWN
P=0.5560
Gene Set Expression (Average CPM)
Pre-Tx Post-Tx
GSEA
NES
FDR
0.980
0.497
e
Paulson et al. 2018
PTIS DOWN
Merkel Cell Carcinoma
No Tx
vs
Avelumab
MCPyV-T cell
Radiation
3.74%
3.79%
tSNE2
tSNE1
High
Avg Log2CPM
Low
Clustered Enrichment Analysis
Gene Set Expression (Average Log2CPM)
All Cells
P=0.6032
Tumor Cells
(CD45-)
****
Macrophages
(CD45+ CD68+)
****
T-Cells
(CD45- CD3D+)
Not Expressed
No Tx Avelumab

**<u>Figure S2</u>: Analysis of $PTIS^{DOWN}$ in published preclinical and clinical datasets after PD-L1 inhibitor treatment**

**(a)** Generation of an αPD-L1 treatment-induced secretome done ($PTIS^{DOWN}$) comprised 4 upregulated genes identified from transcriptome and proteomic analysis using EMT6-PDR cells.

**(b-e)** $PTIS^{DOWN}$ expression in published bulk and single cell RNAseq datasets involving αPD-L1 treatment in preclinical and clinical studies. RENCA-PDR model from this study included.

**(b-c)** Preclinical studies: $PTIS^{DOWN}$ expression using average CPM expression and GSEA in datasets taken from tumor models involving αPD-L1 treatment and found to be **B)** treatment-*sensitive* (GEO: GSE130472, GSE93017, GSE107801) or **C)** treatment-*insensitive* (GEO: GSE130472; RENCA-PDR). Data is compared to vehicle/IgG-treated controls.

**(d-e)** Clinical studies: $PTIS^{DOWN}$ expression using average CPM expression and GSEA in datasets taken from tumor biopsies of αPD-L1 treatment-sensitive patients.

**(d)** NSCLC patients (dbGAP # phs001464.v1.p1): bulk RNAseq from Pre-Tx and Post-Tx tumor sample comparisons (Gray lines indicate matched Pre- and Post-Tx samples).

**(e)** MCC patients (GEO: GSE118056): single-cell RNAseq from untreated (No-Tx) or treated (avelumab) tumor samples with Tsne plots (left) representing average $log_2$CPM expression of PTIS in whole dataset, and bar graphs (right) representing clustered enrichment analysis populations identified by markers for tumors (CD45-), macrophages (CD45+CD68+), and T cells (CD45+CD3D+). Tumor sample that received No-Tx was compared to treated.

*αPD-L1 Treatment-Induced Secretome Down ($PTIS^{DOWN}$); αPD-L1 Drug Resistant (PDR);Counts per million (CPM); Gene set enrichment analysis (GSEA); False Discovery Rate (FDR); Gene Expression Omnibus (GEO); GEO Series records (GSE); database of Genotypes and Phenotypes (dbGaP); t-distributed stochastic neighbor embedding (tsne); Treatment (Tx); non-small cell lung carcinoma (NSCLC); merkel cell carcinoma (MCC).* [#]Indicates genes not part of the Interferome database. *Significance represented as * p<0.05, ** p<0.01, *** p<0.001, **** p<0.0001. Bolded numbers for GSEA represent FDR<0.25 (see Methods).*

**Figure S3**

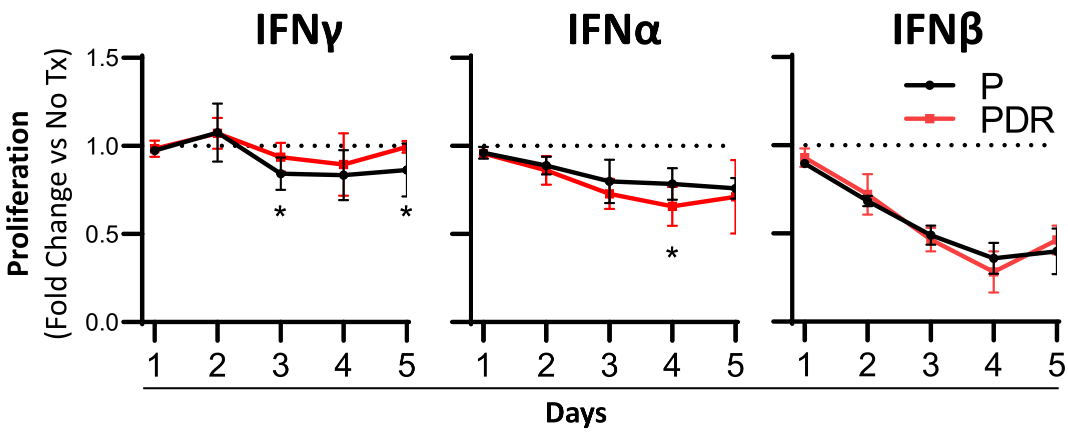


**Figure S3**: ***In vitro* proliferation of EMT6-P and PDR cells after IFN signaling stimulation**

EMT6-P and PDR were treated with IFNγ (left), IFNα (middle), and IFNβ (right) for 5 days and proliferation measured daily by MTS.

*Parental (P); PD-L1 Drug Resistant (PDR); Cells were treated with 10ng/ml of IFNs starting on day 0 and treatment and fresh media was replaced on day 3. * $p<0.05$.*

### Figure S4

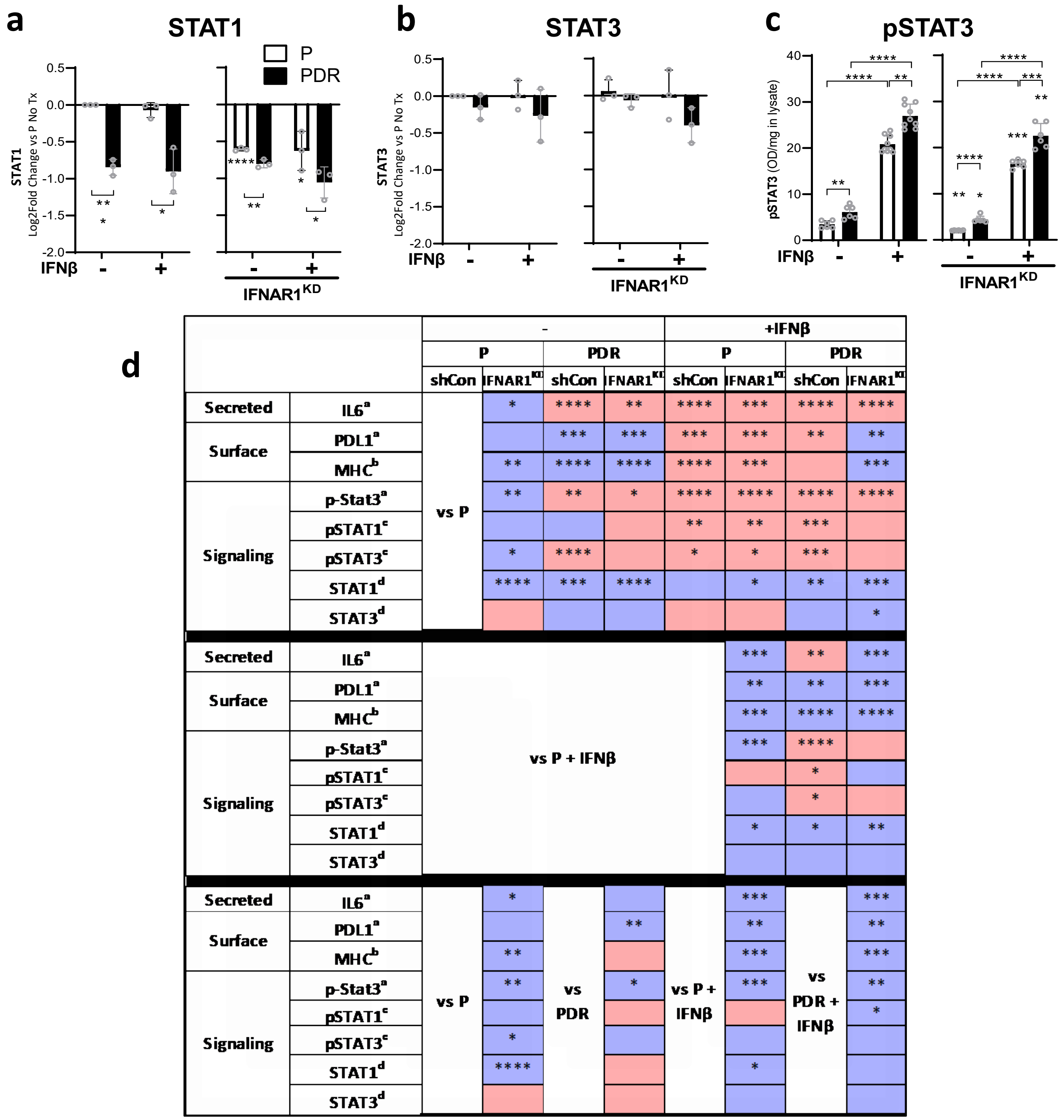


| | | - P shCon | - P IFNAR1KD | - PDR shCon | - PDR IFNAR1KD | +IFNβ P shCon | +IFNβ P IFNAR1KD | +IFNβ PDR shCon | +IFNβ PDR IFNAR1KD |
|---|---|---|---|---|---|---|---|---|---|
| Secreted | IL6[a] | vs P | * | **** | ** | **** | *** | **** | **** |
| Surface | PDL1[a] | | | *** | *** | *** | *** | ** | ** |
| | MHC[b] | | ** | **** | **** | **** | *** | | *** |
| Signaling | p-Stat3[a] | | ** | ** | * | **** | **** | **** | **** |
| | pSTAT1[c] | | | | | ** | ** | *** | |
| | pSTAT3[c] | | * | **** | | * | * | *** | |
| | STAT1[d] | | **** | *** | **** | | * | ** | *** |
| | STAT3[d] | | | | | | | | * |
| Secreted | IL6[a] | vs P + IFNβ | | | | | *** | ** | *** |
| Surface | PDL1[a] | | | | | | ** | ** | *** |
| | MHC[b] | | | | | | *** | **** | **** |
| Signaling | p-Stat3[a] | | | | | | *** | **** | |
| | pSTAT1[c] | | | | | | | * | |
| | pSTAT3[c] | | | | | | | * | |
| | STAT1[d] | | | | | | * | * | ** |
| | STAT3[d] | | | | | | | | |
| Secreted | IL6[a] | vs P | * | vs PDR | | vs P + IFNβ | *** | vs PDR + IFNβ | *** |
| Surface | PDL1[a] | | | | ** | | ** | | ** |
| | MHC[b] | | ** | | | | *** | | *** |
| Signaling | p-Stat3[a] | | ** | | * | | *** | | ** |
| | pSTAT1[c] | | | | | | | | * |
| | pSTAT3[c] | | * | | | | | | |
| | STAT1[d] | | **** | | | | * | | |
| | STAT3[d] | | | | | | | | |

*a, ELISA*
*b, Flow cytometry*
*c, Western comparing phosphorylated levels to total levels*
*d, Western comparing total to β-Actin levels*

Increase
Decrease

**<u>Figure S4</u>: Extended Data from Fig 3. - Protein analysis of ISGs in PDR cells**

**(a-b)** Densitometry quantification of western blots shown in **(Fig 3f)** representing **(a)** relative total STAT1 **(b)** total STAT3 levels compared to total β-Actin levels.

**(c)** pSTAT3 expression in lysates of EMT6-P and -PDR before and after knockdown of IFNAR1$^{KD}$, and after IFNβ stimulation (ELISA).

**(d)** Table summarizing statistical comparisons for data shown in Fig 3 and S4. *Superscripts: a- ELISA, b- Flow cytometry, c- western comparing phosphorylated levels to total levels, d- western comparing total to β-Actin levels.*

*Parental (P); PD-L1 Drug Resistant (PDR); Conditioned Media (CM); Mean Fluorescent Intensity (MFI); IFN stimulated genes (ISGs); IFNAR1 knockdown (IFNAR1$^{KD}$); shRNA vector control (shCon); Cells were treated with 10ng/ml of IFNs and collected after 15mins for STAT1/3 westerns and ELISA. * $p<0.05$, ** $p<0.01$, *** $p<0.001$, **** $p<0.0001$ indicate significance compared untreated controls unless otherwise shown (lines).*

**Figure S5**

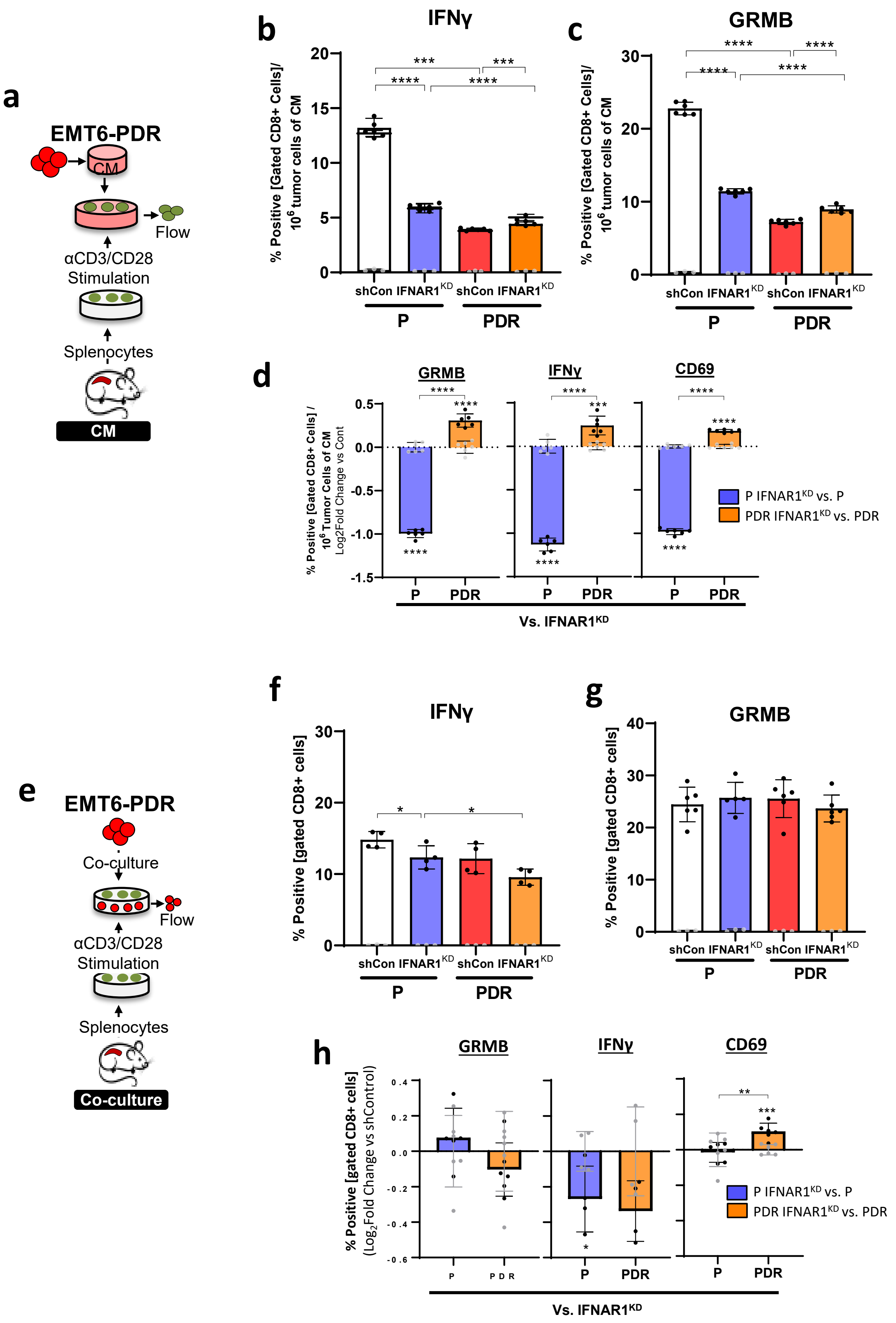
a
EMT6-PDR
CM
Flow
αCD3/CD28
Stimulation
Splenocytes
CM
b
IFNγ
% Positive [Gated CD8+ Cells]/
10^6 tumor cells of CM
shCon IFNAR1^KD shCon IFNAR1^KD
P
PDR
c
GRMB
d
GRMB
IFNγ
CD69
% Positive [Gated CD8+ Cells] /
10^6 Tumor Cells of CM
Log2Fold Change vs Cont
P IFNAR1^KD vs. P
PDR IFNAR1^KD vs. PDR
Vs. IFNAR1^KD
e
EMT6-PDR
Co-culture
Flow
αCD3/CD28
Stimulation
Splenocytes
Co-culture
f
IFNγ
% Positive [gated CD8+ cells]
g
GRMB
h
% Positive [gated CD8+ cells]
(Log2Fold Change vs shControl)
Vs. IFNAR1^KD

**<u>Figure S5</u>: Extended Data from Fig 4. - Splenocyte activation following incubation with PDR cells and conditioned media.**

**(a)** Schematic of Balb/c–derived splenocyte proliferation and activation following incubation EMT6-P and –PDR CM for experiments in b-e.

**(b-c)** CD8+ splenocyte activation marker **(b)** CD69 and **(c)** Granzyme B after co-incubation with CM derived from EMT6-P and –PDR control and respective IFNAR1$^{KD}$ variants.

**(d)** $Log_2$ fold change analysis of CD8+ splenocyte activation markers (Granzyme B, IFNγ, CD69) after co-incubation with EMT6-P and –PDR-IFNAR1$^{KD}$ variant CM compared to respective controls.

**(e)** Schematic of Balb/c–derived splenocyte proliferation and activation following incubation EMT6-P and –PDR CM for experiments in g-j.

**(f-g)** CD8+ splenocyte activation marker **(g)** CD69 and **(h)** Granzyme B after co-culture with EMT6-P and –PDR control and respective IFNAR1$^{KD}$ variants.

**(h)** $Log_2$ fold change analysis of CD8+ splenocyte activation markers (Granzyme B, IFNγ, CD69) after co-culture with EMT6-P and –PDR-IFNAR1$^{KD}$ variant compared to respective controls.

*Parental (P); PD-L1 Drug Resistant (PDR); IFNAR1 knockdown (IFNAR1$^{KD}$); Conditioned Media (CM) * $p<0.05$, ** $p<0.01$, *** $p<0.001$, **** $p<0.0001$ compared to vector controls unless noted otherwise. For C, F, H, and L white bars represent vector controls*

### Figure S6

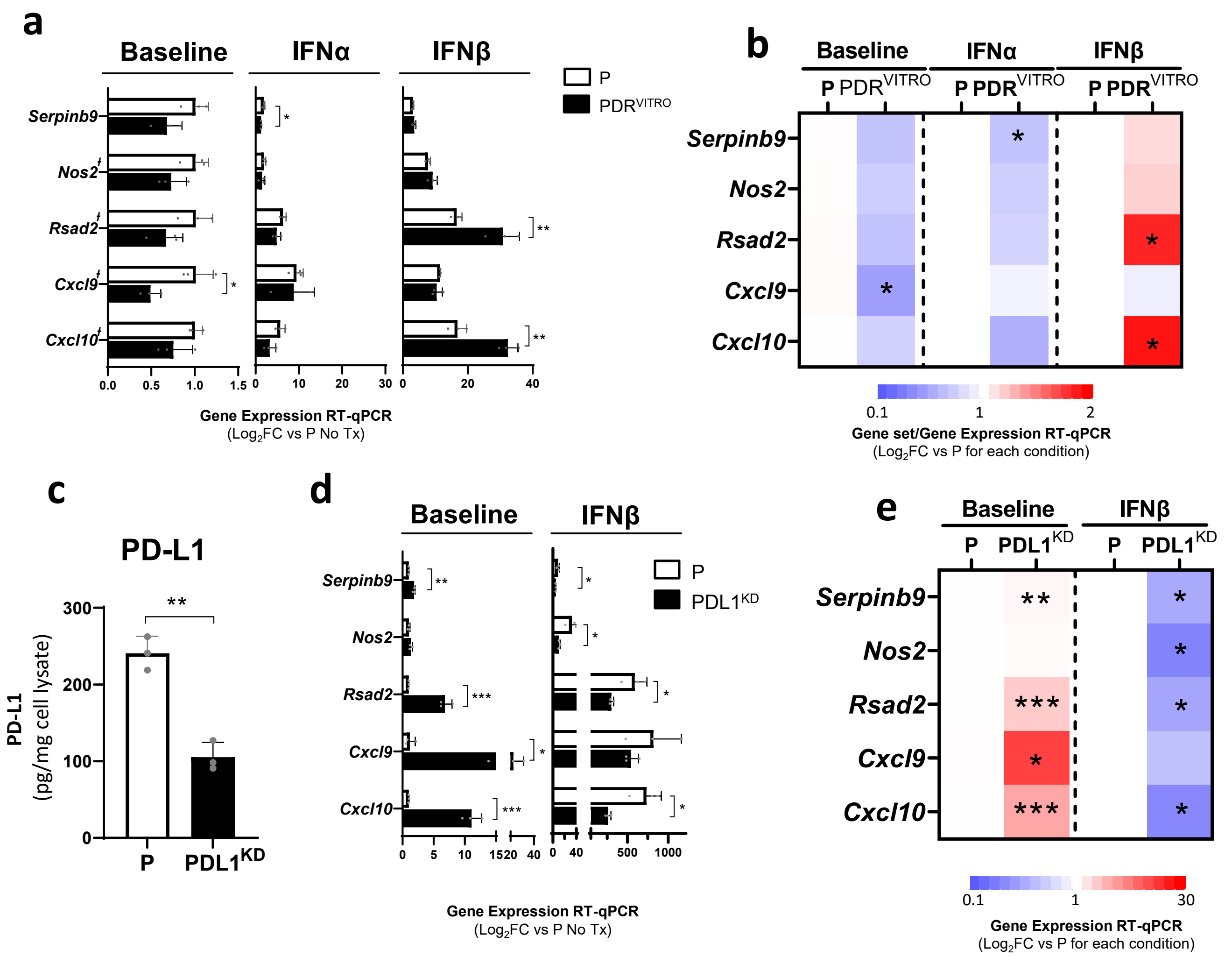


**Figure S6: Extended data from Fig 5- PTIS expression in PDR$^{VITRO}$ and PD-L1$^{KD}$ models**

**(a-b)** PTIS/ISG after stimulation with type I IFNs in EMT6-PDR$^{VITRO}$ cells shown as relative to untreated P controls (a) represented as bar graphs and (b) as heatmap (qRT-PCR). ). ┼ represent genes associated with secretory proteins.

**(c)** PD-L1 expression in cell lysates of EMT6-P and EMT6–PD-L1$^{KD}$ cells (ELISA).

**(d-e)** PTIS/ISG after stimulation with IFNβ in EMT6–PD-L1$^{KD}$ cells shown as relative to untreated P controls **(d)** represented as bar graphs and **(e)** as heatmap (qRT-PCR).

*Parental (P); PD-L1 Drug Resistant (PDR); PD-L1 knockdown (PDL1$^{KD}$) * $p<0.05$, ** $p<0.01$, *** $p<0.001$, **** $p<0.0001$ compared to vector controls unless noted otherwise. For C, F, H, and L white bars represent vector controls*

### Supplementary Tables:

Table S1: Published and Hallmark gene sets used for Gene Set Enrichment Analysis (GSEA)

Table S2: List of PTIS and PTIS$^{DOWN}$ Genes

Table S3: List of Primers Used for qRT-PCR Assays

**Table S1**

| | Gene Sets | Number of Genes | Systematic Name or PMID | Link |
|---|---|---|---|---|
| Published | Liu et al 2018 | 38 | 30559422 | https://pubmed.ncbi.nlm.nih.gov/30559422/ |
| | Benci et al. 2019 ISG.RS | 38 | 31398344 | https://pubmed.ncbi.nlm.nih.gov/31398344/ |
| | Benci et al. 2019 IFNG.GS | 176 | 31398344 | https://pubmed.ncbi.nlm.nih.gov/31398344/ |
| | Weichselbaum et al. 2008 | 49 | 19001271 | https://pubmed.ncbi.nlm.nih.gov/19001271/ |
| | Thorsson et al. 2018 | 24 | 29628290 | https://pubmed.ncbi.nlm.nih.gov/29628290/ |
| | Higgs et al. 2018 | 21 | 29716923 | https://pubmed.ncbi.nlm.nih.gov/29716923/ |
| GSEA MSigDB | Hallmark IFNA | 97 | M5911 | https://www.gsea-msigdb.org/gsea/msigdb/cards/HALLMARK_INTERFERON_ALPHA_RESPONSE |
| | Hallmark IFNγ | 200 | M5913 | https://www.gsea-msigdb.org/gsea/msigdb/cards/HALLMARK_INTERFERON_GAMMA_RESPONSE |

**Glossary**: Molecular Signatures Database (MSigDB), Interferon stimulated genes resistance signature (ISG.RS), Interferon gamma hallmark gene set (IFNG.GS), Interferon alpha (IFNA), Interferon gamma (IFNγ)

**Table S1: Published and Hallmark gene sets used for Gene Set Enrichment Analysis (GSEA)**

#### Table S2

| Anti-PD-L1 Treatment Induced Secretome (PTIS) | | | | | | |
|---|---|---|---|---|---|---|
| PTIS (preliminary) | | | PTIS (enriched) | PTIS$^{DOWN}$ (preliminary) | | PTIS$^{DOWN}$ (enriched) |
| NOS2 | KAZALD1 | PRSS35 | IL6 | CKM | LDLR | ICAM1 |
| LCN2 | SERPINA3N | LUM | CCL2 | MCPT8 | CFP | CSF1 |
| ANGPT1 | GGT1 | CDNF | CXCL10 | MMP10 | S100A3 | COL18A1 |
| COL10A1 | CFH | ISLR | NOS2 | COL4A5 | SRPX2 | HAVCR1 |
| FAM180A | ADAMTS15 | WNT6 | CCL5 | LGR6 | ISM1 | |
| WFDC1 | SNED1 | LBP | CXCL9 | AREG | GPX7 | |
| LGALS7 | GPC6 | SEMA6A | IFNB1 | ADAMTS16 | TNC | |
| ITIH2 | LRRC17 | SEMA6C | LCN2 | MUC16 | PRSS46 | |
| MGP | ELN | C2 | IGFBP6 | CFD | CDCA8 | |
| COMP | ART5 | TNFAIP6 | IL1RN | PF4 | GFRA1 | |
| CLEC3B | OGN | TRIL | CCL20 | WNT5A | ULK4 | |
| INHBE | COLQ | ADAMTS1 | SERPINB9 | OBSCN | SERPINE2 | |
| COL11A1 | CILP | HTRA3 | | TFPI2 | MMP12 | |
| IL33 | KITL | VWF | | CES2E | CXCL14 | |
| APOD | GHR | MMP2 | | COL5A3 | CCL12 | |
| SPON2 | PI15 | ADNP | | FGF21 | GDF15 | |
| FMOD | GBP10 | ADAMTSL5 | | S100A7A | C1QTNF2 | |
| ITGBL1 | C8G | FGFBP1 | | CES1G | LRRN2 | |
| EFEMP1 | GDF3 | PGLYRP1 | | ERFE | TGFB2 | |
| THSD7A | LEPR | F3 | | TMPRSS11B | CPXM2 | |
| CXCL13 | LAMC2 | EDN2 | | CCL22 | S100A11 | |
| OMD | COL3A1 | LGALS4 | | COL17A1 | FAM83D | |
| PRTN3 | HBB-BS | ARSG | | EREG | PODNL1 | |
| 2610528A11RIK | HPN | C1RL | | SFN | OTOS | |
| AGER | SELP | DPYSL3 | | SERPINB2 | PRC1 | |
| S100B | HBA-A1 | TNFSF10 | | CCL11 | COL15A1 | |
| AMY1 | HHIP | POSTN | | IL1RL1 | WNT11 | |
| CRISPLD1 | MATN4 | NXPE4 | | GREM1 | FGF1 | |
| DCN | SERPINA1B | LAMB2 | | PENK | ICAM1 | |
| SEMA3E | HBA-A2 | SAA3 | | HBEGF | CSF1 | |
| ALPL | FAP | ECM2 | | LCAT | HAVCR1 | |
| SECTM1B | GSTM7 | C1S2 | | CEP55 | COL18A1 | |
| LGI4 | TNXB | CD163 | | CCL7 | | |
| MMRN2 | FBLN7 | CCL5 | | CSF2 | | |
| ARSI | HBB-BT | CXCL9 | | MFAP5 | | |
| PRELP | HSPB6 | IFNB1 | | MFAP2 | | |
| PKNOX2 | SEMA3G | CXCL10 | | FREM1 | | |
| R3HDML | MATN3 | IGFBP6 | | GREM2 | | |
| F5 | SLC2A4 | IL1RN | | FABP5 | | |
| XPNPEP2 | OLFML3 | AREG | | MMP9 | | |
| SERPINB1A | LOX | CCL20 | | MMP23 | | |
| EGF | SERPING1 | VEGFA | | PCOLCE2 | | |
| IGF1 | WNT5B | CXCL16 | | MASP2 | | |
| ASPN | C4B | CCL2 | | TMPRSS11F | | |
| | | IL6 | | | | |
| | | SERPINB9 | | | | |

**Table S2: List of PTIS and PTIS$^{DOWN}$ Genes**

**Table S3**

| Gene | Forward Primer | | Reverse Primer | | Mature Amplicon Length |
|---|---|---|---|---|---|
| | Sequence | Length | Sequence | Length | |
| Ifnb1 | GAGCAGAGATCTTCAGGAAC | 20 | AGATTCACTACCAGTCCCAG | 20 | 150 |
| Ifna1 | ACCTTCCTCAGACTCATAACC | 21 | GGGCATCCACCTTCTCC | 17 | 132 |
| Nos2 | GGAGATCAATGTGGCTGTG | 19 | CGGTACTCATTCTGCATGTG | 20 | 108 |
| Il6 | AGCCAGAGTCCTTCAGAG | 18 | GGTCCTTAGCCACTCCTTC | 19 | 148 |
| Cxcl10 | CAGCACCATGAACCCAAG | 18 | TGGCCCTCATTCTCACTG | 18 | 140 |
| Rsad2 | CCCTCTGTGAGCATAGTGAG | 20 | GCCAATCAGAGCATTAACCTG | 21 | 129 |
| Cxcl9 | CCGCTGTTCTTTTCCTCTTG | 20 | GGTCTTTGAGGGATTTGTAGTG | 22 | 135 |
| Serpinb9 | GGCATCAACCATTTAACAAAGAG | 23 | GGCGAGGTTATATGTGTCTTC | 21 | 110 |
| Ccl5 | CCCTCACCATCATCCTCAC | 19 | GTAGAAATACTCCTTGACGTGG | 22 | 137 |

**Table S3: List of Primers Used for qRT-PCR Assays**
