## Supplementary figures and images for "Acquired resistance to PD-L1 inhibition is associated with an enhanced type I IFN-stimulated secretory program in tumor cells"

### Appendix

**Appendix 1**: Western blotting replicates as uncropped unaltered images (related to Figure 3f-g)

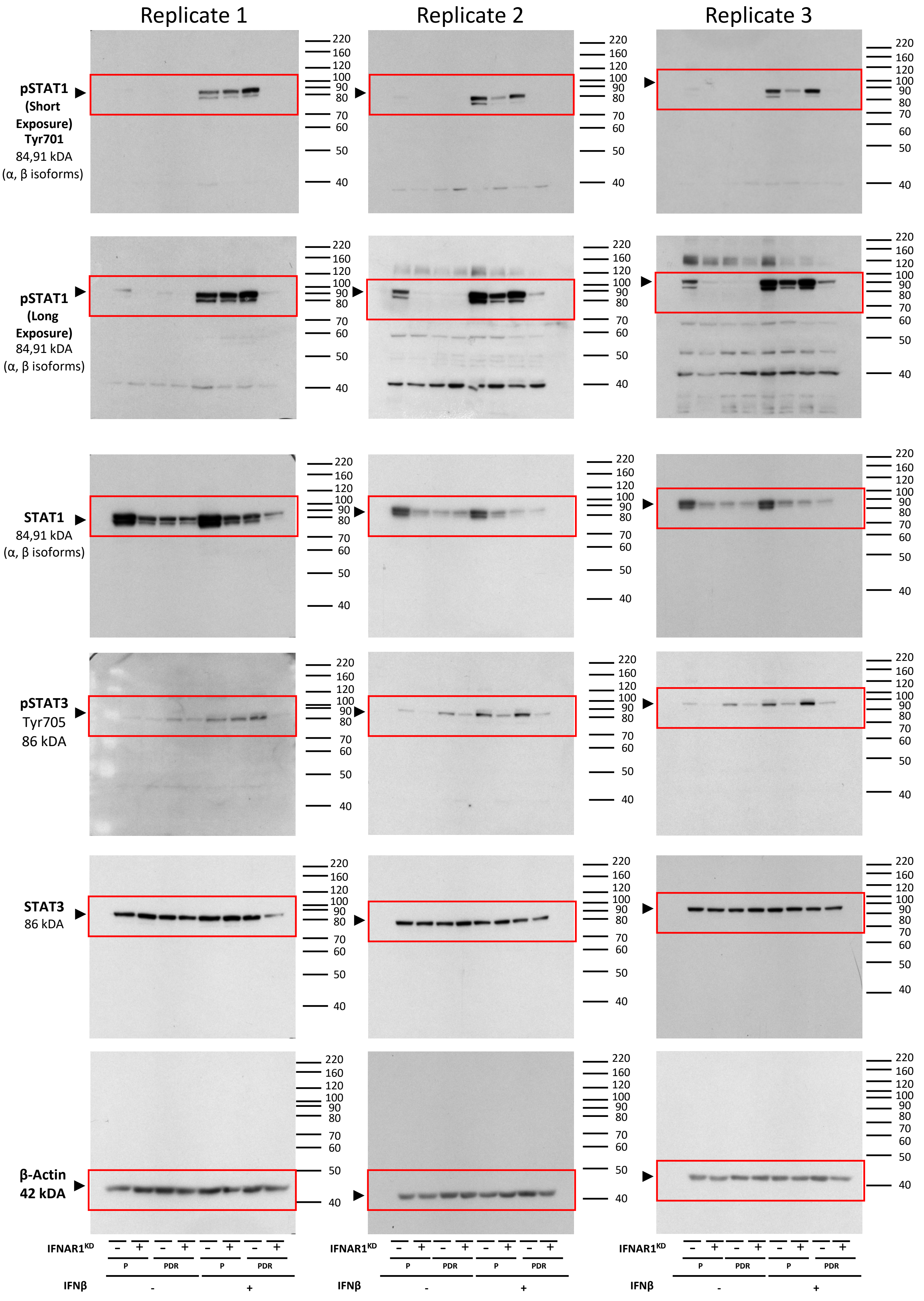
